# Multidimensional mutational phenotypes of MMR deficiency in human cancer cells

**DOI:** 10.64898/2026.07.07.736851

**Authors:** Marcel McCullough, Adam Poti, Maia Munteanu, Fran Supek

**Author notes:** contributed equally (Marcel McCullough, Adam Poti).

## Abstract

Mismatch repair deficiency (MMRd) is a central driver of cancer genome evolution, yet it is commonly reduced to a binary MSI-high versus microsatellite-stable call. Here we combine multiplex CRISPR/Cas12a editing, long-term mutation accumulation experiments in human cells, and whole-genome sequencing to map the mutational consequences of single and pairwise perturbations across canonical and accessory MMR genes, together with selected base excision repair and direct-repair factors. Across 72 isogenic lineages representing 58 genotypes, we recover reproducible substitution, indel and short-tandem-repeat mutational states that distinguish MSH2, MSH6, MLH1, PMS2, MSH3 and MLH3 deficiencies. A dedicated STR spectrum reveals genome-wide evidence for MSH3-driven EMAST and enables design of compact loci panels for mechanism-aware MSI classification. Experimental spectra generalize across independent cell systems and to cancer whole genome sequences, where Elastic-net classifiers accurately identify causal MMR genes in Lynch tumors and uncover common occurrences of intermediate MMRd states. Pairwise perturbations further reveal repair crosstalk reflected in mutational spectra, including enhancement of MUTYH-associated oxidative mutagenesis by MMR loss. Together, these findings recast mismatch repair deficiency as a multidimensional genomic phenotype and provide a framework for interpreting DNA repair deficiencies by bridging mutational phenotypes from experimental systems and human cancers.

## Introduction

DNA mismatch repair is one of the central systems that protects proliferating cells from genome instability. By correcting base-base mismatches and insertion-deletion loops that arise during DNA replication, it preserves sequence fidelity across cell divisions and restrains the accumulation of mutations that can initiate or accelerate tumorigenesis ^1–5^. When mismatch repair fails, the consequences are profound: inherited defects underlie Lynch syndrome and constitutional mismatch repair deficiency, while somatic inactivation is a common route to hypermutation in sporadic cancers. In both settings, mismatch repair deficiency (MMRd) has become biologically and clinically important because it is linked to high mutational burden, microsatellite instability, tumor evolution, and response to immune checkpoint blockade (reviewed in ^6–9)^.

Yet despite this importance, mismatch repair failure is still commonly treated as a relatively simple state in clinical interpretation -- MMR-deficient versus proficient, MSI-high versus MSS. That simplification is not straightforward to reconcile with various individual reports of an EMAST state (reviewed in ^10,11^), which stands for “elevated microsatellite alterations at selected tetranucleotide repeats” and is therefore distinct from MSI-H, which is defined at mononucleotide repeats and dinucleotide repeats. Similarly, the MSI-low (MSI-L) observation in MSI assays may be ascribed to stochastic noise and/or assay artifact ^12,13^, however a possibility remains that it results from bona fide MMRd with different underlying mechanisms ^14,15^. The unidimensional view of MMRd is also somewhat at odds with the accumulating genomic data. Large-scale cancer sequencing, together with emerging experimental systems, has revealed at least seven, and possibly up to nine, different mutational signatures of single base substitutions (SBS) that associate with microsatellite instability ^16–18^. One possible explanation for this is that MMRd is a multifaceted entity; different MMR genes, different branches of post-replicative repair, and different mutational contexts could in principle generate distinguishable patterns of substitutions. Indeed, as a motivating example for this thinking, two out of the seven MMRd signatures have been associated with simultaneous alterations in replicative DNA polymerases ^16,19^. Genome-wide analyses of indels and repeat mutations similarly support that the mutational consequences of MMR loss are richer than the conventional MSI framework implies, and may extend to additional tumor samples and tissue types ^20–24^. At the same time, the field still lacks a clear causal map of these MMRd genomic states.

Mutational signatures have made it possible to infer the action of endogenous and exogenous processes directly from tumor genome sequences, but for DNA repair defects they remain very difficult to interpret mechanistically. In tumors, the observed spectrum is a composite outcome of replication error, endogenous DNA damage, DNA polymerase proofreading, damage tolerance, tissue-specific chromatin and replication programs, clonal selection, and, in many cases, varied mutagen exposures. As a result, a mutational pattern associated with MMR deficiency in cancers may reflect directly the loss of mismatch repair activity, a co-occurring alteration of another DNA repair pathway, a cell type or state that effects DNA repair regulators or cofactors (e.g. free nucleotide pools), or some combination of the three. This interaction is acute for the relationship between mismatch repair and base excision repair, where both pathways can act on overlapping lesions, including deamination-derived mismatches and oxidative damage ^2,5,25,26^. While tumor sequencing has been extraordinarily informative in elucidating the diversity of DNA repair phenotypes, tumors alone cannot fully resolve which genomic patterns correspond to biologically distinct forms of MMR failure as they widely differ in genetic background and tumor cells have undergone various, largely unknown, exposures. Controlled systems are necessary.

Experimental studies of genome-wide mutagenesis in human cells have become tractable and were performed on core MMR genes ^2,27,28^. As challenges, the available systems focus on the mutational outputs of core MMR proteins, consider only single perturbations, apply shorter timescales thereby limiting numbers of mutations for powered statistical analyses, and restrict to a single cell model thus one genetic background. However, DNA repair is inherently buffered and combinatorial. Redundancy within the MutS and MutL families within MMR (reviewed in ^6,7^), crosstalk between MMR and BER and direct-repair pathways ^5,25^, and selective deployment of damage tolerance pathways ^27,29^ can reshape the consequences of losing any one factor in isolation. In such a setting, single-gene perturbation can be incompletely informative, and can miss how repair architecture is organized; to address this need implementations of pairwise perturbations are gaining traction ^30–32^. To systematically elucidate impact of various MMR components, and their interaction within and across pathways would necessitate a controlled human cell system that can combine defined repair perturbations with long-term mutation accumulation, a WGS-based readout of resulting genome-wide mutation patterns at multiple levels -- substitutions, indels, and repeat instability. Further, statistical methods to make these mutational patterns transferrable to tumor genome WGS ^2,3,17^ need to be devised and calibrated, thereby allowing the controlled experiment mutagenesis data to be used for elucidating prevalence and mechanistic bases of MMRd in cancers.

Here we address this problem using combinatorial CRISPR/Cas12a editing ^33–35^ and long-term mutation accumulation in a diverse panel of isogenic human cells derived from the common K562 model cell line. We built a pairwise perturbation set spanning canonical and accessory MMR genes together with selected BER and direct-repair genes, then used WGS to determine how these perturbations reshape mutational landscapes over time. Rather than treating cell line model genomes and tumor genomes as separate problems, we analyzed them jointly, further integrating over substitutions and indel patterns. We complemented this with a dedicated short tandem repeat (STR) framework to capture microsatellite-localized phenotypes that standard signature models compress, and to design focussed, mechanistically discriminative STR panels. This design allowed us to ask questions difficult to answer from tumor genomics alone: whether common MMR genes generate distinct genomic states, whether accessory MMR genes contribute to protection from mutagenesis, whether selected BER or direct repair-related perturbations reshape those states, and whether genome-wide mutational patterns can classify MMRd more richly than conventional MSI-H versus MSS categories.

## Results

### Choice of gene panel spanning MMR and other DNA repair genes

Of the human MMR pathway, four proteins stand out (MSH2, MSH6, MLH1 and PMS2) for their relevance in cancer biology and the known mutagenic phenotype that arises when they are lost. These proteins arrange in heterodimers, forming MutSα (MSH2 and MSH6) and MutLα (MLH1 and PMS2), the core components for the MMR pathway. To unravel the mutagenic role of the different MMR subunits we targeted these four genes and additionally included MSH3, which also heterodimerizes with MSH2 to form the MutSβ subunit, substituting for MSH6. Similarly, out of the core MMR components, germline mutations in PMS2 are the least associated with cancer risk in mouse and human, suggesting that this subunit in the MutLα effector can be partially substituted, thus we included the *MLH3* gene. Furthermore, many MSI tumors lack at least one of these factors due to germline variant, mutation, LOH and/or hypermethylation, yet some MSI tumors appear proficient for these factors ^36^, prompting us to expand the list of MMR-associated genes to test.

Although canonical MMR operates during mitotic replication, related MutS and MutL complexes also function in meiosis, where MutSγ (MSH4-MSH5) and MutLγ (MLH1-MLH3) are required for crossover formation. This motivated us to test if selected meiotic MMR homologs, the MSH5 and also above-mentioned MLH3, also leave detectable somatic mutational phenotypes. Similarly, we also tested SHPRH, as an MMR regulator whose loss has been associated with an MSI phenotype ^36^. Since we aimed to create stable cell lines, we excluded from our list several factors with roles in MMR, such as EXO1, whose fitness effect (via Depmap) we deemed to risk long-term viability of our double KO clones.

To determine whether MMR-associated mutational states are reshaped by repair crosstalk, we paired this MMR-centered gene module with selected genes from base excision repair (BER) and direct repair pathway. These included *MUTYH* and *OGG1* to interrogate overlap between MMR and oxidative-lesion processing, *MBD4* to test a possible interaction with deamination-associated repair, *NEIL2* and *POLB* to sample additional BER activities linked to transcription-associated or gap-filling repair, and *MGMT* and *ALKBH3* to represent direct reversal of alkylation damage. Rather than aiming to exhaustively survey BER, we wanted to test DNA damage classes for which prior biochemical or cancer-genomic studies suggest possibilities of mechanistic convergence with MMR.

### Devising a combinatorial gene knockout system based on CRISPR/Cas12

We modelled MMR and BER gene deficiency using homozygous gene knockout, implemented via an engineered Cas12a (enCas12a) system ^34^. This was better suited than Cas9 for our design because its intrinsic crRNA processing enables compact multiplex guide arrays and therefore efficient pairwise gene targeting in human cells. This feature was key for the present study, as our aim was not only to compare single-gene MMR defects, but also to test selected combinations within and beyond the MMR pathway. We therefore designed dual-guide constructs to generate single- and double-knockout lineages in a common experimental framework.

We built an enCas12a library in which each construct encoded a four-crRNA cassette, with two crRNAs targeting one gene and two targeting a second gene, using modified direct-repeat sequences ^33^. Guides were selected to target separate exons, typically in the first half of the gene, to maximize disruptive editing and increase the likelihood of nonsense-mediated mRNA decay. In our library design the crRNA combinations are not prearranged, instead selected modules can be recovered from a shared oligonucleotide pool via PCR with a certain primer combination optimized for retrieval from mixtures ^37,38^. Next, the Cas12 crRNA modules are assembled into the desired pairwise combinatorial library between the desired modules, building upon a described Golden Gate cloning strategy ^39^, yielding crRNA combinations for all single and all pairwise combinations of genes in the chosen set. Sequencing of the final libraries showed high cloning fidelity (Methods). We then introduced this library into a parental K562 clone stably expressing enCas12a, and recovered individual genotypes by single-cell sorting and subsequent clone expansion **[Fig 1A]**.

**Figure 1.**
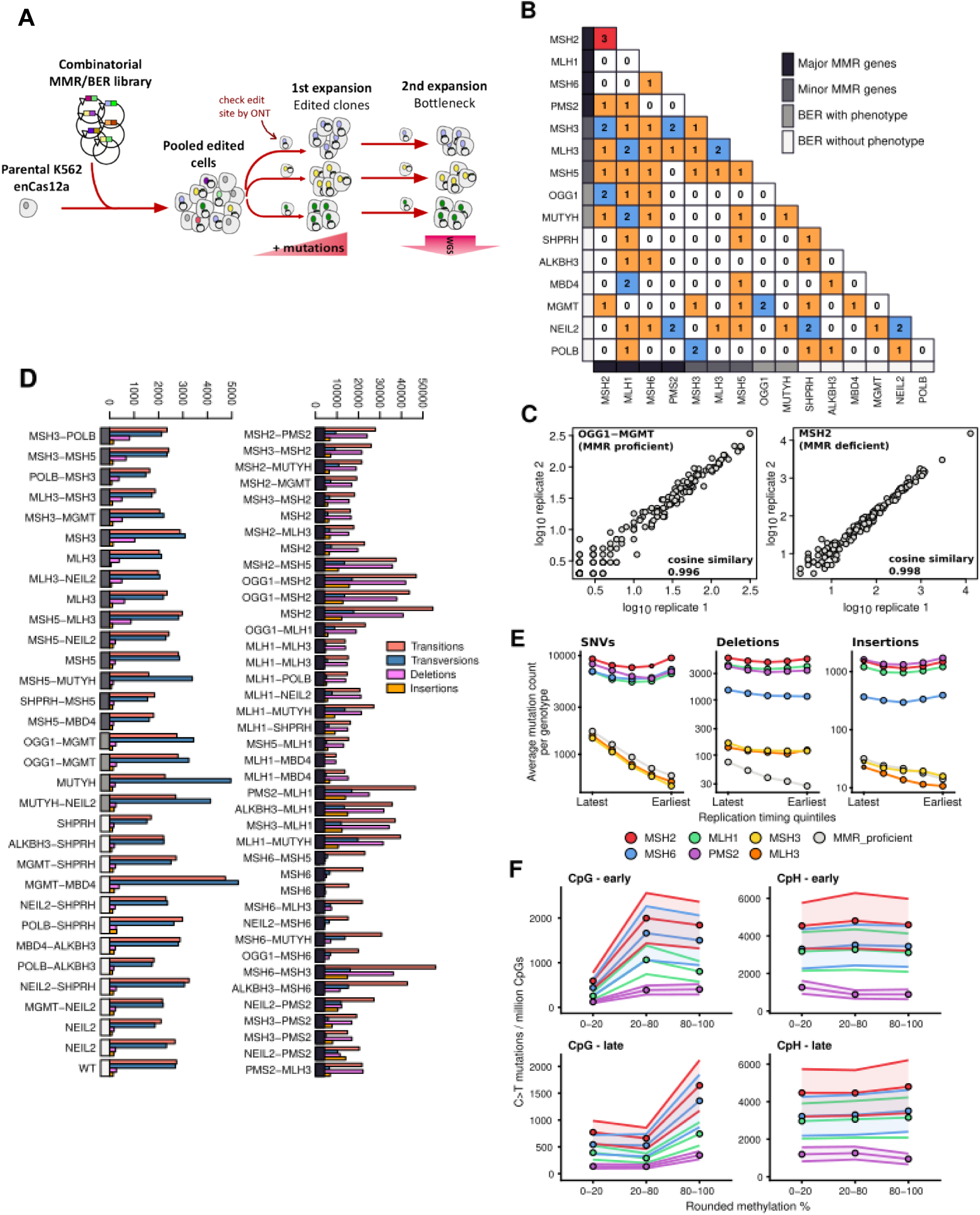
Combinatorial K562 lineages elucidate structured MMR-deficient mutagenesis. **A.** Experimental workflow to generate the combinatorial KOs in a pooled setting, isolate clones and determine genotypes, mutation accumulation, and final bottlenecking before WGS. **B.** Counts of recovered K562 clonal lineages with the indicated gene knock-out combinations and with WGS available. **C.** Consistency of SBS and ID mutational spectra for two example BERd (left) and MMRd (right) genotypes that had biological replicates; remainder in Supplementary Fig. S3. **D.** Mutation counts for SNV transversions, SNV transitions, deletions, and insertions for each of the K562 clones. **E.** Association of mutations with DNA replication timing in K562 clones with the indicated MMRd genotype. **F.** Association of C>T SNV counts with cytosine methylation percentages, shown for MSH2, MSH6, MLH1 and PMS2 cells, grouped by replication timing (early or late) and 3’ nucleotide context (in CpG, or CpH, where H = not G).

Disruptive CRISPR edits on the K562 clones were validated through amplicon Nanopore sequencing. This allowed direct inspection of indels across alleles and provided a practical route to identifying fully-edited clones in this hypotriploid cell line. Dual-targeting (using 2 distinct crRNAs against each targeted gene) improved editing performance: while individual crRNAs showed an average editing efficiency of 73.2%, using two crRNAs per gene increased recovery of full knockout genotypes to 89.9% for single-gene perturbations and 79.8% for double knockouts. Fully edited clones were expanded under physiological oxygen (∼5% O2) for an extended period (multiple months, variable across clones, see **Table S1**) to allow spontaneous mutation accumulation. After that each lineage was bottlenecked again from a single cell before DNA extraction and WGS so that newly acquired mutations would be clonally fixed, and callable with high confidence.

### Mutation accumulation upon combinatorial knock-out of DNA repair genes

We isolated altogether 72 K562 lineages belonging to 58 different genotype combinations, and after whole genome sequencing of each sample (average 28x coverage **[Table S1]**), we called point mutations and short indels by Strelka (joint calling over all samples, removing recurrent variants to exclude germline variation in K562; see Methods). Our mutation accumulation experiment yielded altogether 1,564,693 SNV and 1,057,452 indel mutations **[Fig S1, Fig S2]**, with the least mutated clone (a SHPRH genotype) bearing 3,226 SNV and 290 indel mutations and the most mutated (an example MSH2 genotype) 72,447 SNVs and 52,992 indels. As expected, knock-outs in the canonical Lynch syndrome MMR genes *MLH1*, *PMS2*, *MSH2* or *MSH6* resulted in a marked increase in mutation burden across transition and transversion SNVs and indels **[Fig 1D]**, while other genotypes were associated with more limited effects, e.g., an increase of transversion SNVs in knock-outs in the MUTYH gene in the base excision repair (BER) pathway, or elevation of deletion counts in non-Lynch MMR gene *MSH3* and *MLH3* knock-out clones. We did not detect increased mutation burdens over the *wild-type* clone when considering the BER genes *ALKBH3*, *NEIL2*, *POLB*, the repair regulator *SHPRH*, and the direct-repair gene *MGMT*, nor were their mutational spectra substantially different **[Fig 1D**, **Fig 2A]**. Therefore these knock-outs were not mutagenic in our experimental system and we henceforth consider them effectively neutral, i.e. not substantially alter the mutational signature of a “dominant”, mutagenic MMR or BER gene knocked out in the same clone. In order to test the reproducibility of the mutational signatures across mutation accumulation experiments, we ascertained the similarity between the SBS96 and ID83 mutational spectra of clones with the same k.o. genotype. We found that the cosine similarities are high for all genotypes with multiple mutation accumulation experiments (median = 0.996, range 0.991 - 0.999) and this held true for genotypes equally with low or high mutation counts **[Fig 1B**, **Fig 1C, Fig S3]**.

**Figure 2.**
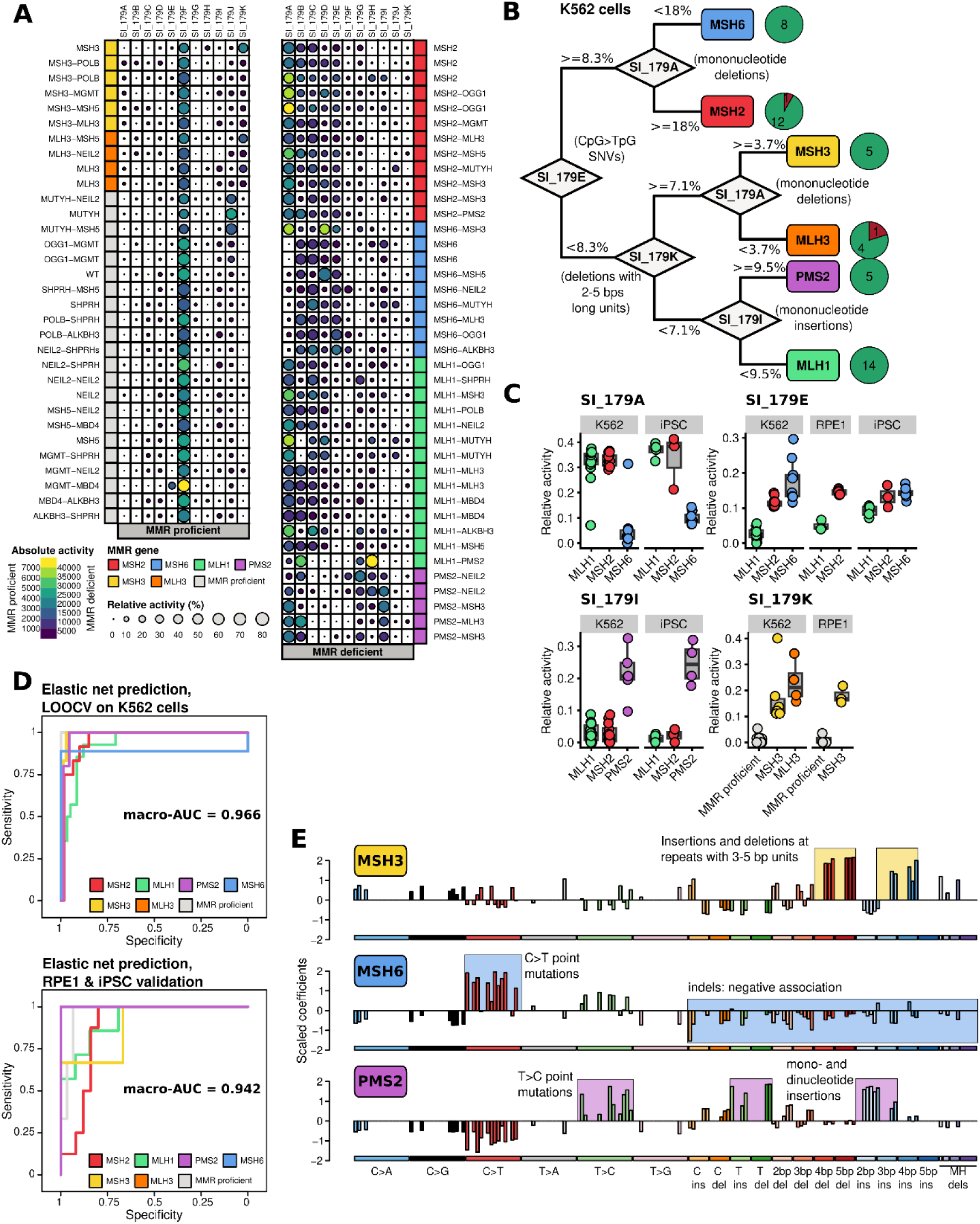
SNV and indel mutational frequencies, merged into a SBS+ID (SI) 179 spectrum, differentially associated with MMR genotypes. **A.** NMF mutational signature exposures in WGS of each K562 clone. Circle sizes refer to relative, circle colors refer to absolute signature activities. **B.** A decision tree separates the major MMR genotypes in the cell line panel. **C.** Refit of K562-based NMF signatures on external in vitro datasets. **D.** Elastic Net model for MMR genotype prediction, assessed via ROC curves and the overall macro-AUC values (average AUC across genotypes). Upper panel: leave-one-out cross-validation on our K562 panel data. Lower panel: testing predictions on independent datasets, with WGS of clones from RPE1 cell type 53 and iPS cells ^2^ (bootstrap with 1000 iterations). **E**. Elastic net coefficients for the prediction of selected genotypes of interest, MSH3, MSH6, and PMS2 genotypes, with the salient feature groups highlighted.

### Distinct mutational states resolve human MMR genotypes

We performed non-negative matrix factorisation (NMF) to determine the main mutagenic patterns in the dataset. We used the concatenated SBS96 and ID83 spectra **[Table S4, S5]** for NMF, as suggested earlier ^40^, henceforth the “SI179” spectrum, and employed the standard SigProfiler tool for *de novo* signature extraction **[Table S6, S7]**. To be able to detect mutagenic processes unique to small groups of samples, we opted for a permissive NMF rank of 11 signatures **[Fig S4, Fig S5B, Fig S6]**. Several components showed an association with specific genotypes **[Fig 2A, Fig S5]** and high cosine similarities to COSMIC SBS and ID signatures **[Fig S7, Fig S8A]**, for example the background signature SI_179F, corresponding to clock-like COSMIC signatures SBS5, SBS40 and ID5, detected in all clones and contributing the majority (73.2 +/− 10.8%) of the exposure in the wild-type clone and in the non-mutagenic BER/DR gene backgrounds. We also recovered the strong enrichment of the signature SI_179J in our *MUTYH* knock-outs; the spectrum of this signature resembles COSMIC SBS36, associated with MUTYH-deficiencies in cancer genomes ^41^.

The stereotyped MMR-deficiency mutational pattern could be observed in MLH1 mutant clones, which could be decomposed into NMF signatures SI_179A and SI_179C [**Fig 2A**], former dominated by COSMIC ID2-like mononucleotide A/T deletions, and latter by a COSMIC SBS44-like SNV component [**Fig S5**]. The widespread presence of the SBS44-like SI_179C across diverse MMR genotypes [Fig 2A] supports the notion of SBS44 being the “basic” MMRd-dependent substitution mutagenic pattern ^18,42^. Three additional NMF components were sufficient to separate the other MMR genes in the K562 panel: our SI_179E, H (or alternatively, the indel-dominant I) and K **[Fig 2B]**. Specifically, we found that components of the MutSα complex (MSH2 and MSH6) can be differentiated by SI_179E, which is characterized by the elevated ratio of CpG>TpG point mutations; our data from the controlled experimental system is consistent with previous reports from observational analyses of cancer genomes ^25,43^. At the same time, SI_179A was more active in *MSH2* than in *MSH6* clones. We mechanistically coupled two distinctive phenotypes to *PMS2* deficiency: SI_179I and SI_179H, representing A/T insertions in repeats and COSMIC SBS26-like T>C SNVs **[Fig S5**], respectively. Finally, *MSH3* and *MLH3* clones could be discerned by a less prominent elevation in SNV and indel counts compared to Lynch MMR genes. These genotypes showed the characteristic SI_179K, exhibiting a relatively high ratio of 2-5 bps long repetitive DNA deletions compared to the more MMRd stereotypical mononucleotide deletions, and having a high similarity to COSMIC ID4 **[Fig S8A, S8B]**, a yet-unannotated indel mutational signature. The mutagenic effects of the MLH3 genotype were observed at high levels in 4 independent clones [**Fig 2A, S1, S2**], with genotypes *MLH3* (2 clones with single gene KO), *MLH3-NEIL2* and *MLH3-MSH5*, implicating the MLH3 protein component of the MutLγ subunit in somatic MMR -- at least in our myeloid cells -- and not only in meiosis as is its widely-recognized role ^44,45^. Thus, MLH3 can likely function *in lieu* of PMS2 and thereby participates in a subset of somatic repair events that appears enriched for loop-type substrates, partly overlapping with MSH3-dependent repair. This surprising finding is supported by biochemical activity of MutLγ on mismatches and loops in cell extracts or using purified proteins, and cooperation with MutSß (*MSH2-MSH3*) was noted ^46,47^. Together, the implication of the above NMF analyses is that SNV (SBS) and indel mutational spectra are largely sufficient to distinguish between the deficiencies in specific MMR genes, at least in this K562 cancer cell model system. This highlights an opportunity to attempt to extrapolate the patterns to other models, and ultimately to cancer genomes to identify specific MMR deficiencies from mutational signatures in the cancer.

### Changes in regional and local mutation risk upon MMR disruption

We further assessed distribution of mutation rates of WGS in individual MMRd genotypes with respect to DNA replication timing (RT), and with respect to DNA methylation, both factors previously associated with particular mutation risk changes upon MMRd in tumor genomes ^25,26,43,48–50^. The stereotypical association of higher mutation rates with later replicating regions ^51^ is seen in the MMRp cell line, and it is abolished in the KOs in all Lynch genes -- MSH2, MSH6, MLH1 and PMS2 -- and this was the case as for SNVs also for indels **[Fig 1E]**, replicating the original observation in MSI-H tumors ^48^ and showing it is largely genotype-independent. We do note that PMS2 has a subtly less pronounced change for SNVs than the other 3 Lynch genes **[Fig 1E]**, consistent with PMS2 mutants retaining residual MMR activity (see above). The MSH3 and MLH3 mutants do not show loss of RT-association for SNVs but they do for indels, specifically deletions (**[Fig 1E]**; insertions are, intriguingly, less affected).

Cytosine methylation is mutagenic by at least two mechanisms: deamination outside of replication, and miscopying during replication. Both potentially result in C>T changes at methylated sites, predominantly in an NCG context in most human somatic cells. These two mechanisms can be influenced by MMR deficiency, however in different ways. First, the deamination lesions (T:G mismatches) may be detected by the MutSα complex and then facilitate BER-mediated repair outside of replication, although the extent of the mutagenic footprint of this mechanism is less clear ^25,43^. Second, the replication errors at methylated cytosines ^52^ would be affected by either MutS or MutL deficiency. We use our isogenic panel of different MMR mutants to solidify this proposed mechanism using data from a controlled experimental system, and to quantify the two effects. Comparing MSH6 mutant (as MutS representative) with MLH1 mutant (as MutL representative), indeed at highly methylated CpG there is approximately double the C>T mutation count in the *MSH6* mutant than the *MLH1* (**[Fig 1F]**, top left), while in the control CpH context (H = not G) the C>T rates are near identical in *MSH6* and *MLH1* (**[Fig 1F]**, top right). This distinction is observed to a similar extent in the early-RT and in the late-RT regions (**[Fig 1F]**, top row vs bottom row), suggesting it largely stems from a replication-independent process. Consistently with DNA methylation being the cause, the difference in CpG mutagenesis between *MSH6* and *MLH1* diminishes in lower-methylated regions (**[Fig 1F]**; that it does not disappear fully might be explained by somewhat mismatched DNA methylation between our long-term cell culture and the reference methylation track), while this is not the case for the case of co-located CpH dinucleotides. The effect is qualitatively similar if instead of *MSH6*, *MSH2* is considered as a MutS representative, again normalizing CpG (**[Fig 1F]** left) with CpH control context (**[Fig 1F]** right). Overall, our data supports that it is a mutagenically significant mechanism that MutS specifically (without MutL involvement) deals with CpG lesions, presumably deamination-induced mismatches, largely outside of DNA replication, handing them over to another DNA repair pathway; we could not confirm specifically the role of MBD4 ^25^ (not shown). Overall, by assessing the MutS-KO to MutL-KO mutation rate differences, we suggest that a roughly similar amount of C>T mutations at methylated CpGs are prevented by this MutS-specific, BER cooperation mechanism, as the C>T mutations that are prevented by the MutS-MutL replicative mismatch correction.

### A robust, generalizable genome-based classification of MMR deficiency mechanisms

After identifying the broad mutational patterns in our data that included MMR deficient genotypes, we set out to generate a predictive model that draws on individual components of the mutation spectrum to classify MMR deficiency mechanisms, with the outlook towards application to cancer genomes. To this end, we used a multinomial Elastic net regression framework, trained on the mutations from our K562 isogenic cell line panel, and validated in 3 external datasets (see below). As features, we utilized the relative mutational spectra, the total mutational burden, the SNV/indel ratio and the “age” (here, maintenance time) of the individual clones. Assessed by leave-one-out crossvalidation, the Elastic net models was able to efficiently differentiate the individual MMR genotypes (average AUC across classes = 0.966) **[Fig 2D, upper panel]**, with the lowest AUC=0.889 for the MSH6 mutant genotype (n=9 WGS in training data, which includes one apparently erroneous prediction that results from a genetic interaction in MSH6-MSH3 double KO; see below for details; excluding that instance, the MSH6 AUC=1).

We validated the Elastic net models for individual MMRd mechanisms on the two external datasets with mutation accumulation in human cell culture: ^53^ that used isogenic RPE1 cells (retinal epithelium) to generate single KOs in the genes *MLH1*, *MSH2* and *MSH3* that overlapped our gene set, and additionally ^2^ that used iPSCs (stem cells) with single KOs in overlapping genes *MLH1*, *MSH2*, *MSH6* and *PMS2* **[Table S8]**. The remaining mutagenic genes were unique to our study with respect to these external datasets (*MLH3*, *MSH5*, *MUTYH*, *OGG1*) and thus the models predicting these mechanisms could not be validated therein. On these data, the model to sort MMR deficient cells by the gene inactivated reached an average AUC of 0.944 (AUC = 0.902 and 0.984 for RPE1 and iPSC cells, respectively) on the batch effect-corrected data (see Methods) [**Fig 2D, lower panel]**. The coefficients of the Elastic net model served as an independent alternative for NMF signatures for feature importance assessment: we found that the feature effect sizes and directions showed a good correspondence to the above-mentioned feature combinations extracted by NMF. For example, this supported the role of 3-5 bps unit size indel events for MSH3 differentiation, the underrepresentation of indels and the enrichment of C>T SNVs in MSH6 samples, and the importance of T>C SNVs and insertions in PMS2 samples **[Fig 2E]**. The most important predictors for distinguishing MutS and MutL mutant clones were the higher level of NpCpG > NpTpG SNVs in the former group [**Fig S9**].

### Genomic MMR classifier is portable to cancer WGS and identifies correct underlying gene in Lynch patients

To validate that our MMR deficiency mechanism calls transport to cancer genomes, we utilized the presence of Lynch syndrome patients in a TCGA cohort subset (including 182 cases in COAD/READ ie. colorectal cancer, and STAD and UCEC cancer types). Our TCGA subset was selected to enrich for MSI-H and MSI-L cases (Methods), specifically, we analyzed WGS of 60 MSI-H, 84 MSI-L and 38 MSS cases **[Table S9]**. We curated a ground-truth set of MMRd genomes via the presence of Lynch syndrome inferred via MMR gene-inactivating pathogenic germline variants (see Methods) providing a strong prior of the causative gene, and thus a control and validation of our method if it should predict the expected causative MMR gene from the mutational spectra **[Table S10]**. By using batch effect correction (Methods), we were able to align the mutational spectra of TCGA cancer samples and our K562 clones, removing the confounding effects of different mutation burdens (cancers are expected to have undergone more cell divisions) and any cell type-specific differences **[Fig S10A, S10B]**. Additionally, to explicitly model uncertainty, we assigned a quantitative confidence score to each prediction using bootstrapping (Z_def_ score: the delta of top-ranked vs second-ranked genotype probabilities, divided by the bootstrapping confidence interval of the top-ranked probability **[Fig 3A]**; see Methods); this implies that some predictions could be classed as “medium confidence” or assigned to a MMR deficiency type at different thresholds. After making predictions for the 13 Lynch cases from the TCGA dataset and the three MSS control samples with low mutation burdens, we found that in 11 out of 13 cases the model correctly predicted the causative MMR gene, of those in 7 cases with high and 4 cases with low confidence [**Fig 3B**]. Importantly, in the 4 medium-confidence and 2 incorrect cases, the tentative causative genotype was the second most-probable category **[Fig S11]**. The model correctly identified the three MSS cases with high confidence as “MMR proficient” as well.

**Figure 3.**
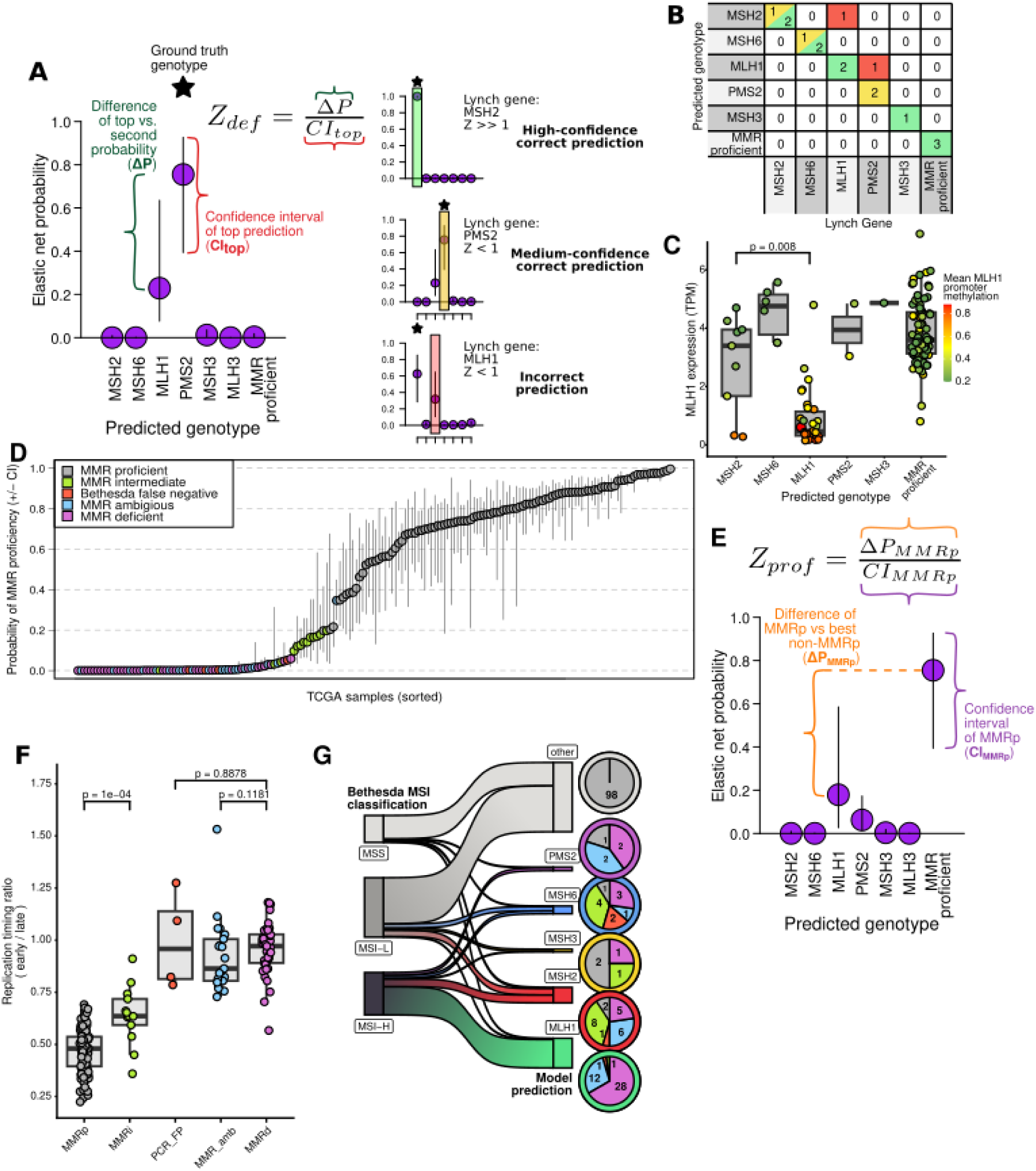
Establishing MMR deficient (MMRd), intermediate (MMRi) and proficient (MMRp) states via an uncertainty-aware classification method. **A.** Schematic of the assessment of the Elastic Net models’ predictions for individual MMRd genotypes and comparison with MMRp state. The Z_def_ score compares the confidence interval (inferred via bootstrapping) of the top-ranked genotype, versus the distance between the top-ranked and the second-ranked genotype Elastic Net probabilities; Z_def_ >= 1 implies a clear distinction between MMR genotypes, while samples with Z_def_ < 1 are labelled as “medium-confidence”. **B**. Predictions of the K562-derived Elastic Net model, evaluated across an external WGS dataset of presumed Lynch cases (carriers of deleterious germline variants in MMR genes) in TCGA cohort. Colors in confusion matrix correspond to three scenarios in A panel. **C.** Associations between the Elastic Net predictions, and epigenetic MLH1 status (assessed by promoter methylation and mRNA levels), only shown for high-confidence genotype predictions. **D.** Sorted probabilities of the MMRp label with confidence intervals across the TCGA sampled set of colorectal, uterus and stomach cancers. Colors denote prediction confidence groups, and conflicts with the Bethesda PCR assessments (see Results). **E.** Schematic of the assessment for identifying MMRi status. The Z_prof_ score estimates the confidence of separation from the MMRp group, and a Z_prof_ < 0.5 indicates that the given sample cannot be reliably differentiated from MMRp. **F.** Association of mutation rates with DNA replication timing (early/late ratios) in MMRp versus 3 MMRd categories (MMRd, high-confidence genotype-specific MMRd; MMR_amb, MMRd with uncertain genotype; and PCR_FP, false positives of the Bethesda PCR assay; see Fig S14), and MMRi as the intermediate case. Individual samples shown in Supplementary Fig S14. **G.** Re-classification of the considered TCGA cases using the MMRd and MMRi rules established herein. Within the pie chart, colors (as in panel D, F) show the most likely confidence-category, for each of the predicted genotypes (the colored rings around the pies; the green is the MLH1-like, red is MSH2-like spectrum etc.).

To assess the robustness of our Elastic net MMRd model to out-of-distribution mutational spectra, we analysed predictions on TCGA WGS cases that exhibited activity of strongly mutagenic processes apart from MMRd, and were not included in the training data. First, we grouped MMR proficient cases according to the activity of SBS17a/b, a signature of uncertain etiology, possibly reflecting incorporation of oxidised nucleotides into DNA and/or misincorporation of uracil ^54,55^. We found that the prediction probabilities of MMR proficiency were very similar between cases with more vs less than 1000 SNVs attributed to SBS17a/b **[Fig S12A, S12B]**. We also assessed predictions for hypermutated TCGA cases caused by SBS10a/b/c/d, mutational signatures linked to pathogenic mutations in the exonuclease domains of replicative DNA polymerases delta and epsilon **[Fig S13A, S13B]**. 11 of the 15 cases with SBS10a-d hypermutation were classified as MMR proficient, implying that the MMR Elastic net classifier largely correctly discarded the markedly divergent mutational spectra in this group, despite being unseen during training on cell line data. Of the 4 remaining SBS10 cases predicted as MMRd, the 3 (with MSH6 phenotype predictions) were marked as ambiguous confidence, and there was 1 PMS2-predicted confidently classified by the model. While the 4 SBS10 cases are, nominally, misclassifications, 2 cases were actually MSI-H annotated by the Bethesda PCR assay and all 4 bear hallmarks of *bona fide* MMRd: a mutational excess in C>T and T>C SNVs (for three MSH6 and one PMS2 case, respectively), and elevated indel/SNV ratios [**Fig S13B**]. Finally, as a complementary analysis we considered the converse case of the SBS17 and SBS10 out-of-distribution spectra assessed above: cancers that have another mutagenic process but it is combined with a likely MMR deficiency, asking if that MMRd would still be detected by our Elastic net model. In particular we assessed tumors with high activity of SBS14 or SBS20 signatures **[Fig S13A]**, associated with DNA polymerase epsilon or delta pathogenic mutations concurrent with MMRd ^19^. Our Elastic net indeed detected MMR deficiency in all 7 of these SBS14/20 dominant, presumably DNA polymerase/MMR double-deficient samples. For 6 out of 7, it was predicted to have the *MSH6* label, in accordance with enriched C>T SNVs and a decreased SNV/indel ratio for this genotype [**Fig S13C**]. Overall, our MMRd classifier based on SBS and ID spectra appears robustly applicable also to tumors with various non-MMR background mutational signatures, exhibiting few false positives.

### Genomic MMR classifier is sensitive and uncovers additional MMRd and “MMR intermediate” tumors

Next, we made predictions of MMRd for the WGS of the relevant cancer types in the TCGA cohort, and compared them to the pre-existing MSI assessments in TCGA metadata, based on the PCR of Bethesda STR loci, which are available for COAD, READ, STAD and UCEC cancer cases [**Table S9**]. As anticipated, the majority of cases annotated as MSI-H by the Bethesda panel were predicted by our method to be deficient for MLH1. This is supported by the cross-reference of model predictions with MLH1 expression and promoter methylation values [**Fig 3C**]: cases predicted to be MLH1-null had significantly lower MLH1 expressions, and higher average promoter methylation values.

Interestingly, we found various TCGA cancer cases (5/38 MSS and 19/84 MSI-L samples; 13% and 23% respectively) diagnosed as MSS or MSI-L by Bethesda panel PCR that were predicted to be MMR deficient by our WGS-based Elastic net model. To estimate confidence in our predictions and thus support or discard the possibility these hits signify a real subgroup, we again used the bootstrap estimates of uncertainty, here to assess separation from MMR proficiency state (Z_prof_: delta between MMRp probability and best-scoring MMRd probability, divided by the confidence interval of MMRp probability **[Fig 3E, Fig S14A]**; see Methods). We sorted all TCGA cases into five categories based on this measure, our Elastic net predictions and SNV/indel ratios [**Fig S14B**]. Cases that were assigned MSS or MSI-L by the Bethesda panel PCR assay and that had non-MMR-proficient Elastic net predictions were divided into two groups: a) not distinguishable from MMRp i.e. likely false positives of our method (Z_prof_ < 0.5; 7/24), and “MMR intermediate” (MMRi) cases with either genome-wide mutational characteristics of MMR deficiencies, e.g. SNV/Indel ratios < 1 (probable false positives of the nomBethesda assay, 4/24), or by confident predictions (Z_prof_ >= 0.5) in our panel (13/24) **[Fig 3D]**. Finally, cases that had a Z_prof_ >= 0.5 and Z_def_ < 1 --meaning they are confidently MMRd, but only with a “medium-confidently” determined which exact genotype-- were classified and “MMR ambiguous” (MMR_amb).

To obtain independent evidence for the occurrence of a partial MMR deficiency in these MMRi cases, with relatively lower mutational burdens, we assessed the ratio of mutations in the early vs. late DNA replication timing (RT) genome domains, a known correlate of MSI-H status ^48^; importantly, this statistic is in principle independent of the trinucleotide and indel mutational signature analysis. Encouragingly, we see a clear correlation between the Elastic Net Z_prof_ scores from SBS and ID mutational signatures, and the late RT association of mutation rates in individual samples (R = 0.76), clearly extending beyond the MSI-H/MSS dichotomy **[Fig S14A]**. Our “PCR/Bethesda false negative” group had identical early-vs-late RT ratios as nominally MSI-H cases, supporting they are strongly MMRd but undetected by the standard clinical assay. Crucially, the more abundant “MMR intermediate” group showed a significantly different, intermediate loss of RT association [**Fig 3F**]. This supports a moderate MMRd phenotype in MMRi group (∼22.6% of nominally MSI-L and ∼13.1% of nominally MSS samples), and solidifies that our Elastic net-based MMR classification pipeline reliably provides a mechanism-grounded cancer patient stratification using WGS global mutational patterns.

### STR mutational spectra of indels accurately classify genotype-specific repeat instability patterns

As microsatellite instability is a hallmark of MMRd, we explored whether a more detailed, fine-grained spectrum of genome-wide patterns of indels at short tandem repeat (STR, synonymous to microsatellite) loci can be derived, yielding a statistical tool for more accurate and more interpretable mechanistic classification of MMR genotypes, compared to the standard ID83 spectrum. Specifically, we generated indel calls for each locus based on allele length histograms generated with the MSISensor tool (see Methods), and categorised them according to a custom, STR-specific classification scheme to generate STR238 mutational spectra **[Fig 4A, Fig S15]**. This scheme used features such as repeat unit length and sequence, repetition length, indel type and length, binary replication timing and Alu element overlaps, comprising 238 channels in total.

**Figure 4.**
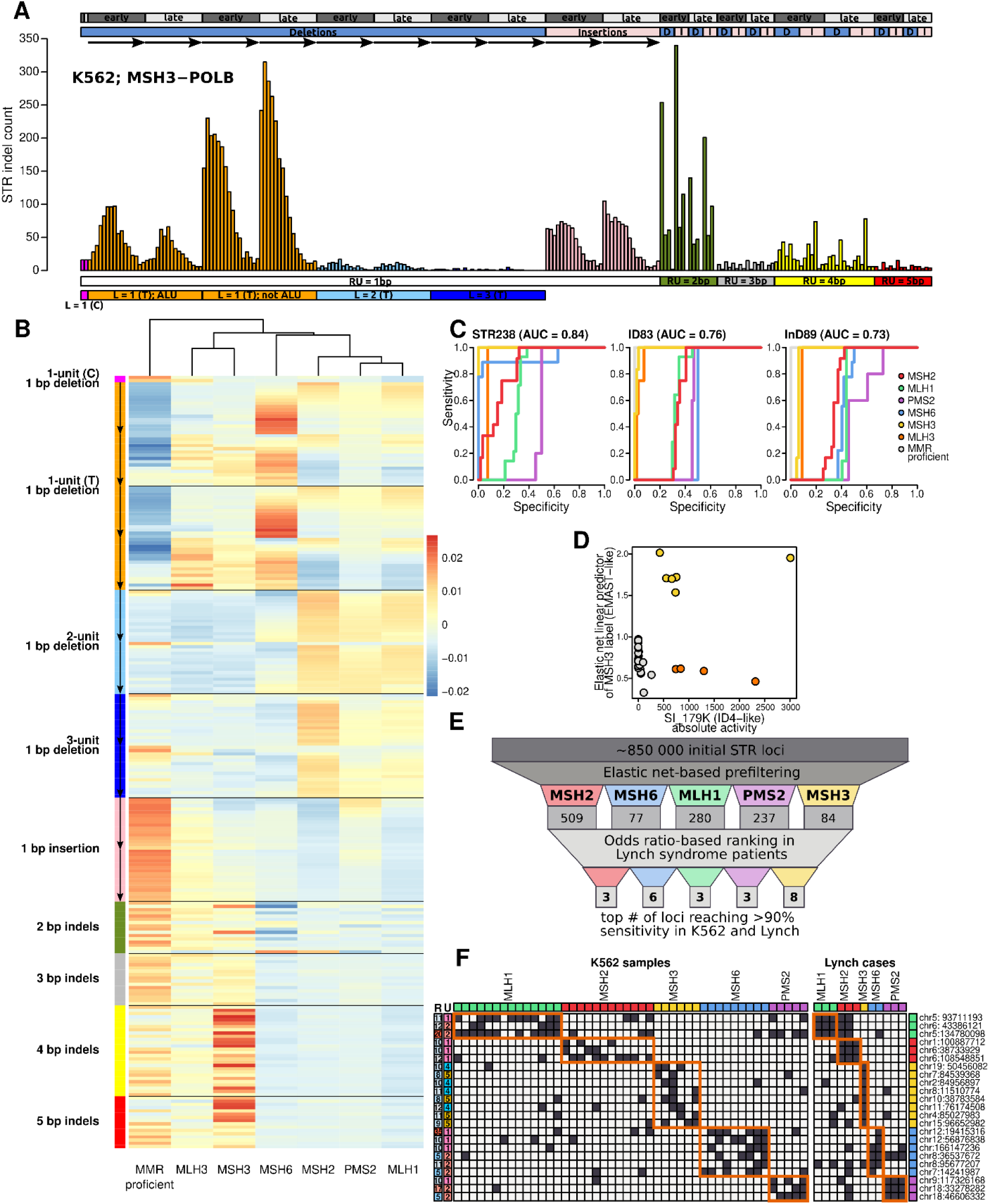
Design of the STR238 spectrum of instability at short tandem repeats and mechanistically-informed genome-wide STR panels. **A.** Example of the design of short tandem repeat spectra, shown for the case of OGG1-MLH1 double KO genotype; RU = repeat unit size, D = deletion, I = insertion, L = deletion/insertion size in repeat units, the black arrow symbolize the gradient of repetition lengths from 10 to 25+. **B.** Elastic net coefficients (scaled) for classifying MMR genotypes in the K562 panel by the STR238 spectrum. The black arrows indicate the gradient of repetition sizes from 10 to 25+ in the case of mononucleotide repeats. **C.** Accuracy of Elastic Net models (as ROC curves) of MMR genotype separation in K562 using three types of indel-only spectra: STR238, compared with previous ID83 and InD89 spectra. **D.** The NMF component STR_sigF is the manifestation of EMAST, and describes EMAST more precisely when contrasted with SI_179K signature (related with COSMIC ID4), since it can separate MSH3 (more plausibly, direct EMAST cause) from indirect effect of MLH3. **E.** Design of a mechanistically-grounded STR locus panel to capture MMR gene-specific instability phenotypes, by synergizing cell line model WGS and tumor WGS data. **F**. Mutational status (black: mutated, white: not mutated) of each selected STR locus in the K562 cells, and selected TCGA Lynch patients, with the relevant genotypes. On the left, shown repetition length “R” and repeat unit length “U” for each locus in the STR panel.

We compared the STR238 spectrum to the standard COSMIC ID83 spectrum, and to the recently reported InD89 spectrum that proposes to better discriminate patterns resulting from defects in DNA replicative polymerases, in their ability to distinguish the MMR deficiencies in our cell line clones, showing a superiority of STR238 to both alternative indel spectra (mean AUC over genotypes = 0.84, compared to 0.76 and 0.73 for ID83 and InD89, respectively **[Fig 4C]**). We note, as per our analysis above, the standard indel ID83 spectrum gains high accuracy in this test if supplemented with the SBS96 spectrum, supporting the notion of a “multimodal” approach to deriving mutational signatures ^40^.

In addition to the STR238 spectrum’s power to dissect mechanisms, its detailed categorization aids interpretation, thus allowing insight to be gained from the pattern [**Fig S16A**]. Similarly to SBS96+ID83 spectra, again in this STR238 spectrum we employed a multinomial Elastic net approach to find the most characteristic STR indel classes for the individual MMR genotypes. [**Fig 4C**]. Additionally, our data supports that longer mononucleotide tracts are mutated at higher rates ^21^, with the rate decaying above repetition lengths of 25 units, possibly determined by technical limits of short-read alignment. In the case of *MSH6* knock-outs, the lengths of repeats harboring mononucleotide repeat deletions were higher (median length of mutated STRs in MSH6: 15.25, versus MSH2: 14.49, MLH1: 14.72), estimated via Elastic net model’s importance for these categories. In accordance with decreased activity of MutSα over MutSß on longer DNA loops, we found a decreased indel rate of *MSH6* knock-outs affecting dinucleotide vs mononucleotide repeat sites [**Fig S16C**], and among mononucleotide repeats, its mutation spectra are also “shifted” towards the more mutable, longer homopolymers. In case of *PMS2* clones, the Elastic net coefficients also imply the distinctive role of one-unit insertions in 10-15 units long mononucleotide STRs **[Fig S16B]**; the phenotypes of MSH2 and MLH1 deficiency are not distinguishable in this STR spectrum. The above illustrates how the various repeat length categories, distinct in our STR spectrum but collapsed together (all beyond 5 nt) in the standard ID83 spectrum, can provide an additional data source informative of the genotype identity of given samples.

As an interesting side finding, we observed that the ratio of 2- and 3-unit deletions **[Fig S17A, S17B]** and the weighted means of the mononucleotide repetition lengths [**Fig S17C**] is dependent on the cell line age (maintenance time during long-term culture in the MMRd state [**Table S1**]), suggesting that various multi-unit deletions have actually resulted from consecutive single-unit deletions.

### Support for EMAST in the STR and conventional ID83 signatures

An additional benefit of STR238 spectrum is in assessing mutagenic processes operating at repeats with longer unit lengths. This signal is not well captured by the COSMIC ID83 spectrum, which collapses together all unit lengths of 2 and longer, meaning trinucleotides and tetranucleotides are not distinguished from dinucleotide repeats. Moreover this is a blind spot for the standard Bethesda panel PCR based method for determining MSI, which covers only mononucleotide and dinucleotide repeats and moreover does not distinguish them. Here the STR238-based classification uncovered an exclusively MSH3-dependent process, also acting on longer unit repeats [**Fig 4C**]. While indels affecting mononucleotide repeats were biased towards deletions in all MMR genotypes, these longer unit indels were more balanced between insertions and deletions [**Fig S16A**]. Especially tetranucleotide repeats were overrepresented compared to MMR proficient and, as a comparator, MLH3 clones, and among these, tetranucleotide STRs with AGAT and AAAT units [**Fig S16D**]. Indel counts of the MSH3 mutant in these categories were comparable to the counts in MSH2 and MLH1 samples, but not MSH6, consistent with the expected distinction of function between MutS subunits [**Fig S16A**] where MutSα (MSH2-MSH6) should not detect loops at tetranucleotides. We note that our analysis on the joint SBS96+ID83 spectrum above (via NMF component SI_179K) did capture this trinucleotide/tetranucleotide repeat signal indirectly, via identifying 2-5 bp deletions in relatively short repeats. However this signal was less precisely delineated, as it was active in both MSH3 and the MLH3 clones, and moreover the differential effect on different unit length STRs would be collapsed such that tetranucleotides could not be identified as the primary target.

These characteristics suggest that we provide here the direct whole-genome evidence of the EMAST phenotype, a mutational process previously suggested to be linked to MSH3 silencing or dysfunction in cancers (reviewed in ^10,11^) and in our data indeed resulting from ablation of the MSH3 gene in a human cell line model. While our SI_179K signature (with an COSMIC ID4-like component) was fairly accurate in distinguishing MSH3 and MLH3, considered jointly, from other MMR genes, the STR spectrum makes it possible to further distinguish between the MLH3 and MSH3 signatures [**Fig 4C**], the MSH3 having more changes specifically of lengths ≥4 [**Fig 4B**]. This underscores the value of STR-based spectra in interpreting mutational signature mechanisms and contextualizing them in prior knowledge, such as here having connected the SBS+ID signature SI_179K with EMAST.

### Design of targeted STR loci panels for detecting mechanism-specific genome instability

The genotype-specific STR mutagenesis patterns suggested that the WGS data we derived in controlled experiment could be used to design a minimal panel of STR loci that would distinguish individual MMRd mechanisms. In a sense, this is a generalization of the established NCI/Bethesda five-locus panel and its unidimensional MSS/MSI-L/MSI-H calls to a multidimensional classification framework that can indicate the exact MMR gene affected. We used our dataset of mutated STR loci in our K562 isogenic cell lines as a starting point, and by a multi-step filtering setup (see Methods) we selected 3-8 repeat sites per genotype (total n=23 loci) whose mutational status showed >90% sensitivity for the genotypes *MSH2*, *MSH6*, *MLH1*, *PMS2* and *MSH3* **[Fig 4D]**. As a validation, we inspected these microsatellites status in WGS of presumed 12 Lynch cases in TCGA with known causal genes, where there was considerable association **[Table S10]** thus many of the loci would be considered replicated across cell types and systems in this proof-of-concept analysis for STR panel design **[Fig 5E]**. Challenges arise with individual loci in distinguishing certain combinations (particularly MSH2 versus MLH1, and in distinguishing MSH6 from PMS2 or MLH1) although in all validation Lynch cases a combination of the 23 loci would be fully predictive of the gene affected. As further support of validity, this STR feature-agnostic approach resulted in STR sets bearing the previously-known STR characteristics per genotype, for example, MSH3-specific loci were tetra- and pentanucleotide repeats **[Fig 5E]**. Overall, we demonstrate the possibility of designing genetic panels -- here, containing microsatellite loci -- for precise inference of mutational mechanisms by drawing on controlled mutagenesis WGS data.

**Figure 5.**
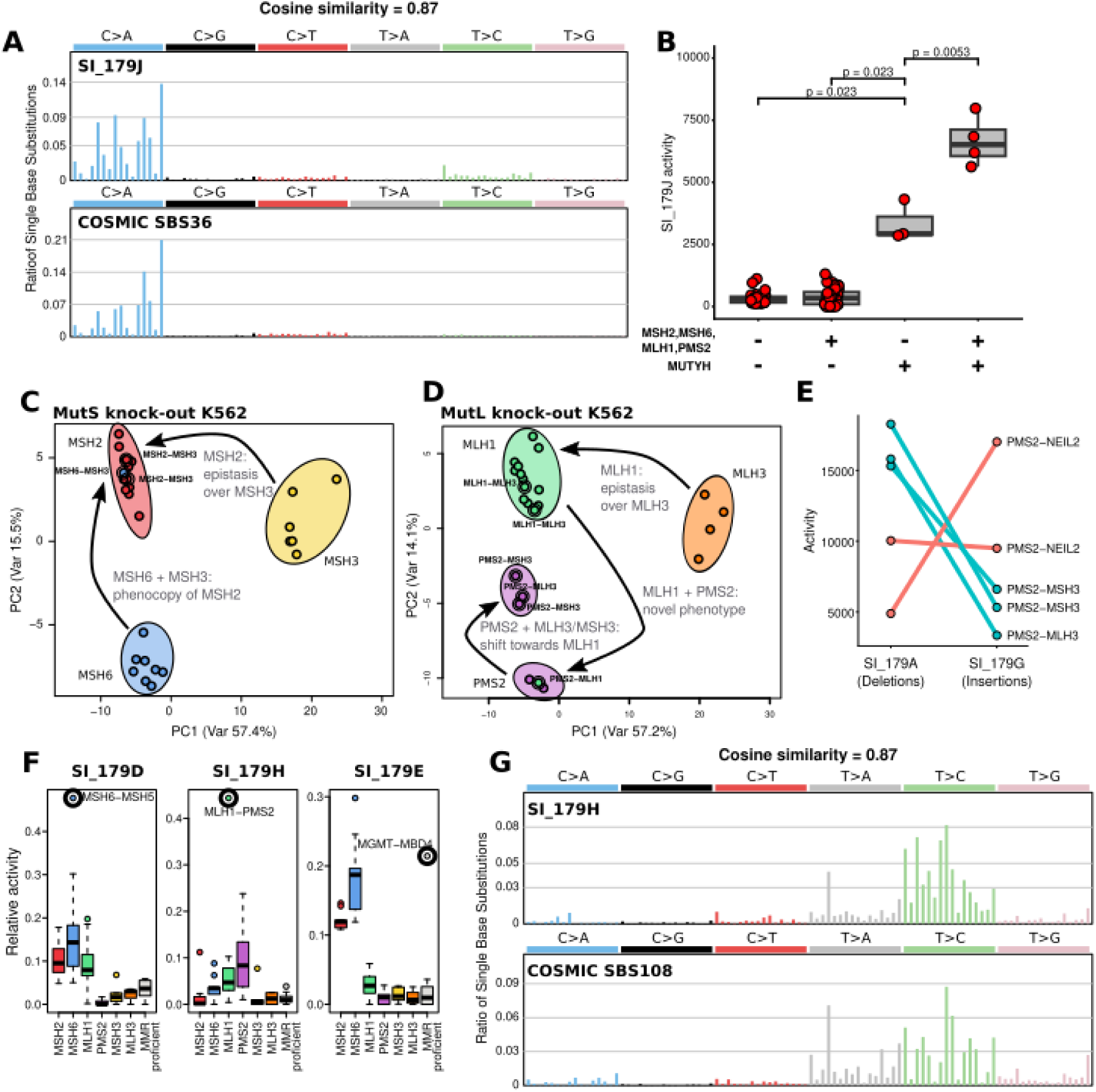
Epistatic interactions within and between MMR and BER pathways assessed by mutational spectra of combinatorially-perturbed human cells. **A.** Comparison of trinucleotide mutational spectra of our SBS+ID signature SI_179J and the COSMIC BERd-associated SBS36. **B.** Activity (“exposure”) of the signature SI_179J in MUTYH single mutant clones, MMR gene (MSH2, MSH6, MLH1, and PMS2) single mutants, and double-mutants. **C.** Principal component analysis (PCA) of SBS+ID (SI_179) spectra in WGS of K562 clones with MutS complex mutations, with indications of MMR double mutants and their position changes in the PC space. **D.** As panel C, but with MutL complex mutations. **E.** Activities of signatures SI_179A (dominated by 1bp repeat deletions) and SI_179B (dominated by 1bp repeat insertions) in PMS2 mutants: concurrent knock-out of MSH3 or MLH3 decreases the insertion/deletion ratio. **F.** Activities of “orphan” (single-clone) mutational signatures (SI_179D, SI_179H and SI_179E) in the K562 isogenic panel; thick circles indicate the clone with dominant contribution. **G.** Comparison of SI_179H, which we found in a single K562 sample with MLH1-PMS2 genotype and COSMIC SBS108, a recently described signature without assigned etiology.

### Combinatorial perturbation reveal epistatic interactions across DNA repair pathways reflected in mutational spectra

Our K562 cell line panel contains pairwise gene perturbations, and is therefore suitable to discover genetic interaction within and across the MMR and BER pathways. We found SI_179J in MUTYH knock-out cells **[Fig 5A]**, and showed that it shows a cosine similarity of 0.87 with COSMIC SBS36, a mutational signature associated with MUTYH deficiencies in human tumors **[Fig 5B]**. Interestingly, we found that the activity (NMF “exposure”) of SI_179J was significantly higher in 2 MLH1-MUTYH, 1 MSH6-MUTYH and 1 MSH2-MUTYH lineages where MUTYH and one of the major MMR genes are concurrently inactivated (1.98-fold increase between means, t-test p = 0.0053, Cohen’s D = 3.79) **[Fig 5A]**. This suggests a substantial role of MMR in countering mutations resulting from oxidative lesion mismatches remaining upon MUTYH loss, where both MutS and MutL components of MMR play an important role.

Next, we noted an elevated relative activity of SI_179E in a single MMR-proficient clone with the genotype of MGMT-MBD4 [**Fig S4**]. SI_179E is a combination of COSMIC SBS1 and SBS44 **[Fig S7]**, and differentiated MSH2 and MSH6 from MLH1 (see above). As SBS44 is the quintessential MMRd-dependent mutagenic pattern, its presence seemed improbable in an MMR proficient sample therefore suggesting this example would be mechanistically distinct. Upon inspection of the mutational spectrum, we noted that CpG>CpT mutations specifically are increased [**Fig S18]**. This SBS1-like admixture is consistent with overlapping function of MGMT and MBD4, for instance demonstrated when dealing with mutagenic O6-methylguanine lesions ^56^, and with the known causal role of germline MBD4 mutations in SNV mutagenesis with a SBS1-like spectrum in human tumors ^57^.

### Pairwise genetic knockouts within MMR pathway suggest interactions and roles for accessory MMR proteins

Our experiment generated 20 lineages of K562 cells where two components of the MMR pathway were inactivated at the same time. This enabled us to analyse the organization of MMR via epistasis in observed mutagenic outcomes. Regarding the MutS complex, here serving as a control, we found that the mutational phenotype of MSH2-MSH3 double mutants is very similar to the general MSH2-specific mutagenic patterns **[Fig 5C]**, in agreement with MSH2 being an obligate partner for MSH3 activity. Similarly, a concurrent loss of MSH6 and MSH3 phenocopies the MSH2 knock-out phenotype, meaning that MSH2 is only functional in complex with either MSH6 or MSH3. These examples illustrate how WGS-derived mutational signatures can be used to detect epistasis in a controlled experimental system, thereby elucidating DNA repair pathways.

In the MutL complex **[Fig 5D]**, a double knock-out of *MLH3* and *MLH1* was identical to the general MLH1 pattern, consistent with MLH3 having MLH1 as an obligate partner in repairing mismatches or DNA lesions. However the MLH1-PMS2 double mutant did, unexpectedly, generate a unique SBS spectrum (discussed below). Moving on to PMS2, the concurrent inactivations of PMS2 and MLH3 modified the characteristic PMS2 phenotype, partially shifting it towards the MLH1 loss (i.e. full MutLα loss) as evident in for instance decreasing the insertion/deletion ratio **[Fig 5E]**. Interestingly, similar was observed for PMS2 and MSH3. Regarding the STR spectrum, we noted a difference in the double mutants: while mono-, di- and trinucleotide repeats showed decreasing deletion and increasing insertion numbers, in tetranucleotide or longer-unit STRs, both deletions and insertions decreased, again underscoring the role of MSH3/MLH3 in EMAST [**Fig S16**]. This implies that (i) MLH1 is functional without PMS2 and MLH3, possibly by binding the PMS1 and (ii) the insertion bias in PMS2 knock-outs is caused by MLH3, likely acting with MSH3, which have been shown previously to act jointly in repeat expansion disorders ^58^.

Finally, we set out to analyse two “orphan” signatures, i.e., NMF components from the joint SBS96+ID83 NMF that were found only in a single double-mutant in the current dataset [**Fig 5F**]. First, signature SI_179D was found in a single MSH6-MSH5 knock-out, can be characterized by a surplus of Y<u>C</u>W>Y<u>T</u>W mutations, and suggests an intriguing possibility of a mitotic role of MSH5, which is normally considered to be active during meiosis ^59^. Our clones also contain MSH5 co-knockouts with other MutS proteins (1 clone of MSH2-MSH5 and MSH3-MSH5 each; Fig 1c) which do not exhibit this mutational signature, suggesting either it is a MSH6 specific effect, or possibly an orthogonal biological occurrence in that clone e.g. secondary mutation in another DNA repair or chromatin modifier gene.

Second, SI_179H was present in a MLH1-PMS2 double mutant lineage (one clone available in our panel), and was enriched in A<u>T</u>R>A<u>C</u>R and C<u>T</u>K>C<u>C</u>K point mutations (where R=A or G, and K=G or T). Interestingly, this pattern showed a cosine similarity of 0.87 with COSMIC SBS108 [**Fig 5G**], a novel addition to the COSMIC signature set, found recently in a subset of MSI colon cancers (see Supplementary Note in reference ^60^), suggesting that these MMRd colon cancers might have a similar mechanistic cause. An outstanding question is, given that mutational phenotype of this clone appears different than of the MLH1-only mutant (with some resemblance to the PMS2 KO, see all spectra in Fig S1), what residual activity allows MMR to proceed when only MLH1 is ablated but PMS2 remains functional, and how is this modified when PMS2 is additionally deleted.

## Discussion

By combining long-term mutation accumulation with defined pairwise perturbations in human cells, we show that mismatch repair deficiency is better understood as a structured landscape of related but distinct failure modes, rather than as a single, binary instability label. MSH6-deficient cells consistently accumulated fewer indels than MSH2/MLH1/PMS2-deficient cells but remained hypermutated by substitutions, while PMS2 deficiency was marked by a distinctive bias towards insertions at repeats and SBS26-like T>C substitutions; these are consistent with a prior experiment ^2^ and with various observations from cells and tumors of patients with MMR deficiencies ^18,24,61–63^. MSH3-deficient cells occupied a lower-burden region of indel mutational phenotype space in our data, mirroring recent perturbation experiments in mouse and human models ^4,53^, and the spectrum was recognizable by longer deletions at repeats. Our methodology provided confidence estimates, robustly distinguished MMR genotypes in independent in vitro datasets, and (after aligning experimental and tumor spectra) the same predictive model transferred to human cancer genomes wherein it correctly classed most tested Lynch cases. Taken together, these results argue in favor of that the observed heterogeneity in mutational signatures observed in MMR-deficient tumors could have resulted from disruptions in different MMR genes and, possibly, combinations thereof.

The mutational states we observe are plausible mechanistically, including ones resulting from genetic interactions. On the MutS side, the fact that MSH2-MSH3 double mutants phenocopied the general MSH2-deficient state, and that combined loss of MSH6 and MSH3 likewise converged on the MSH2-null mutational phenotype (same was observed with cancer risk phenotype in mouse ^64^), strongly supports the validity of our experimental and analytical framework; MSH2 is known to function as the obligate scaffold for the lesion-recognition branches of MMR. At the same time, also in line with known biochemistry, the relative sparing of indels in MSH6-deficient cells is consistent with partial buffering by the MSH3-containing branch for loop repair. On the MutL side, the data argue against PMS2 loss being equivalent to full shut-down of repair; the distinctive PMS2 mutational phenotype with T>C transitions and relative enrichment of insertions ^2^ also observed here, its modification by concurrent loss of MSH3 or MLH3, and the unique spectrum observed in the MLH1-PMS2 double mutant all suggest that residual or rerouted repair remains possible in human somatic cells. This would likely occur by MLH3 partially substituting for PMS2 to form the MutLγ complex, which retained *in vitro* residual repair capacity for base-base mismatches and single-nucleotide loops ^46^, and is supported by mouse models of cancer risk and microsatellite instability combining germline Mlh3 and Pms2 ablation ^65^. Somewhat speculatively, we suggest a somatic role of PMS1 as well, as concurrent loss of PMS2 and MLH3 still did not result in a MutL-null (i.e., MLH1-deficient) mutational phenotype. As an evolutionary contrast, *C. elegans* lacks orthologs for the alternative MutL subunits MLH3 or PMS1, and therein when *pms-2* is knocked out, the resulting mutation rate and mutational profile are identical to *mlh-1* worm mutants ^66^. Mutation burden and signature data in our study strongly support that MLH3 does have roles in human somatic MMR. The mutational signature association with MSH3 is consistent with a proposed MMR axis activity where MSH3 (MutSβ) interacts with and functionally engages the MLH3-MLH1 (MutLγ) complex ^47,67^. Conceptually, we demonstrate how mutational spectra can serve as information-rich functional readouts of repair architecture: they not only report that MMR has failed but also reveal which components remain active, elucidating where compensation is taking place.

Various studies from the pre-genomic era have reported EMAST marked by instability at tetranucleotide loci, and we find evidence of its existence across tumors via the genome-wide SBS+ID mutational spectrum classifier trained on the MSH3-ablated experimental model. Our current estimation is that the MSH3-like mutagenesis occurs in cancer at modest frequencies: about 3% in our mixed-tissue data set, which was enriched for MSI-L. This is in contrast with pregenomic studies that suggested widespread EMAST ^68,69^ and better aligned with a WGS study where ∼3.5% (8 of 224) of MSS colorectal cancers had some enrichment in indel mutations at tetranucleotide repeats relative to total indels, even though a conventional PCR-based fragment analysis would have classified 20% of their MSS tumors (8 of 40 tested) as exhibiting instability at >2 tetranucleotide loci ^70^.

As for the MSI-L phenotype, in contrast to reports based on genomics claiming it is practically equivalent to MSS with respect to absolute indel burden ^13,21^, we do classify a subset of MSI-L tumors as MMRd based on relative mutational spectra. Such MMR-deficient MSI-L have enrichment in MSH6-like mutational signatures, compared to tumors exhibiting MSI-H proper, which were enriched with MLH1-like signatures; consistently, across MMRd colorectal cancers, a lowered expression of MSH2-MSH6 was linked to a weaker mutator phenotype than loss of expression of MLH1-PMS2 ^61^.

Interestingly, we reclassify a minor subset of the nominally MSS tumors as MMRd, and additionally introduce an “MMR intermediate” (MMRi) state based on WGS mutational signatures. Because this reclassification of nominally MSS or MSI-L tumors into MMRi tumors finds independent support in an orthogonal mutational phenotype characteristic of MMR deficiency -- “flattening” of the domain-scale mutation rate landscape in relation with DNA replication time ^48^ -- these predictions appear reliable. MMRi labels suggest that some degree of MMRd are more widespread than the MSI-H phenotype derived by the NCI/Bethesda clinical diagnostic assay, which is recognized to undercall mild or less-common forms of MMR failure e.g. via MSH6 loss ^71,72^, as well as to be variably accurate across tissues ^9,72,73^.

Our study recasts microsatellite instability as a multidimensional phenomenon, via deriving a more resolved STR spectrum that aids interpretation and accurately predicts the underlying MMR gene which was altered. This provides an additional tool for mechanistic dissection from WGS and moreover allows for the design of a compact STR panel with good accuracy for identifying the underlying cause of the MMRd, as we validate on TCGA cancer WGS data.

These STR spectra showed an MSH3-specific process enriched, expectedly, at tetranucleotide repeats thus providing direct genome-scale evidence for EMAST from a controlled experimental system, and (unlike the standard SBS and indel spectra) the STR spectrum of MSH3 was distinguished from MLH3. Another interesting example is that PMS2-deficient samples displayed enrichment of insertions, potentially resulting from a putative, less well characterized DNA repair cooperation between MutSβ (MSH2-MSH3) and MutLγ (MLH1-MLH3) ^67^, since this insertion enrichment disappears when MSH3 or MLH3 are deleted in this PMS2-deficient background. Therefore our findings suggest that MLH3 -- as expected, having a secondary role in somatic DNA repair compared to PMS2 -- drives an atypical MMR pathway that is more prone to insertions.

The combinatorial design of the study also allowed us to probe interactions across DNA repair pathways, in particular between MMR and BER. The interaction between MUTYH loss and core MMR defects stands out, boosting the SBS36-like mutagenesis normally resulting from MUTYH disruption ^2,74,75^ but not altering the spectrum. This interpretation is consistent with that MMR recognizes a broader repertoire of lesions apart from replication mismatches ^5,25^. Our data suggests a comparable increase in MUTYH mutagenesis in 2 MutL clones (MLH1-MUTYH), and in the 2 MutS clones (MSH2-MUTYH and MSH6-MUTYH), therefore arguing for only a minor role for a mechanism where the MutS complex acts as general DNA damage sensor and “hands off” lesions to another repair pathway. Observing this MUTYH interaction, considered together with MMR interactions with DNA polymerase alterations and alkylation damage ^19,27^, raises the possibility that various other signatures seen in tumors might in fact reflect mixed or sequential engagement of MMR with exposures or with deficiencies in other pathways. Some pairwise k.o. states we observed (MSH6-MSH5 and MLH1-PMS2 double mutant) produced a mutational spectrum that was not predictable from either single perturbation. These illustrate how possible novel biology may be revealed via gene combinatorial perturbations and subsequent mutation accumulation experiments, however we highlight the need for replication in independent studies.

Overall, our study supports the paradigm where challenges in assigning mutational signatures to mechanisms are well-addressed by systematic gene perturbation experiments, and we advance that combinatorial gene editing is particularly well-suited to resolving redundant and interacting DNA repair pathways. In the case of MMR, this strategy revealed an elaborate landscape of multiple, overlapping yet mechanistically distinct genomic states that can be recognized in human cancer WGS, with implications to modeling tumor evolution and, potentially, stratifying for prognosis and therapy.

## Data and code availability

The genetic variant calls generated in the study will be made available upon manuscript acceptance. Further, the results published here are in part based upon data generated by the TCGA Research Network [https://www.cancer.gov/tcga] available under restricted-access applications placed via dbGaP [https://www.ncbi.nlm.nih.gov/projects/gap/cgi-bin/study.cgi?study_id=phs000178.v11.p8], and data was downloaded from Genomics Data Commons data portal [https://portal.gdc.cancer.gov/]. Code will be made available upon manuscript acceptance.

## Acknowledgements

A.P. was supported by Juan de la Cierva postdoctoral fellowship from the Spanish government. Group of F.S. was supported by an ERC Starting Grant “HYPER-INSIGHT” (757700), ERC Consolidator Grant “STRUCTOMATIC” (101088342), Horizon2020 RIA project “DECIDER” (965193), Horizon Europe project “LUCIA” (101096473), Spanish government project “REPAIRSCAPE”, CaixaResearch project “POTENT-IMMUNO” (HR22-00402), a Novo Nordisk Fonden “Start Package” grant, the Danish Cancer Society grant “AI-DRIVERS” and a DFF Project2 (5243-00072B), the SGR funding of the Catalan government, and the Severo Ochoa Centers of Excellence award to the IRB Barcelona. LLM tools were used for language editing of manuscript text. We are grateful to group members drs. Marcos Moreno, Aleix Bayona, Sergio Marin, Daniel Naro and Alejandro Modrego for excellent technical assistance and/or scientific discussions.

## Author contributions

M.Mc. co-designed the experiments, performed experimental work (including plasmid design, cloning, transformation, cell culture, gene edit validation, DNA extraction), interpreted data, and participated in drafting the manuscript; A.P. peformed computational analysis (mutational signature analysis, integration of tumor and cell line data, devised statistical methods, performed validation), visualized data, interpreted data, and co-drafted the manuscript; M.Mu. performed bioinformatics analyses of cell line data (including QC, variant calling and filtering), and participated in mutational signature analysis; F.S. co-designed the experiments, interpreted the data, co-drafted the manuscript, and supervised the work.

## Methods

### Cell culture and isolation

HEK293T cells were cultured in DMEM supplemented with 10% fetal bovine serum (FBS), 1% penicillin-streptomycin (P/S). Cells were maintained in a humidified incubator at 37°C with 5% CO2. K562 cells were cultured in RPMI-1640 medium, supplemented with 10% FBS and 1% P/S. Cells were maintained in a humidified incubator at 37% and 5% CO2 during transduction and selection steps. After cell sorting, clones were kept in normoxia conditions (5% oxygen) to minimize oxidation-related mutations from being kept at atmospheric conditions (21% oxygen). Cultures were passaged once or twice a week according to their growth rate.

Single-cell sorting was performed using a FACSAria II Flow Cytometer (BD Biosciences) using a 100 μm nozzle tip. The target population was first identified based on forward scatter (FSC) and side scatter (SSC) parameters to distinguish single cells from debris and doublets. Sorted cells were collected in 96-well plates containing 200 μL of supplemented RPMI-1640 medium. The plates were then incubated at 37°C with 5% CO2 and 5% O2 and surveilled for growth.

### crRNA selection and Oligonucleotide pool design

Two crRNA per gene were selected from the top picks generated through the CRISPick algorithm ^39,76^, which has been optimized for enCas12a targeting. The crRNAs were prioritized by highest predicted efficiency, low off-target activity and had to target the beginning of the gene (excluding the first exon). Same gene crRNA target separate exons, when possible, and not have more than 3 consecutive thymines, nor have the BsmBI endonuclease recognition site (CGTCTC) nor its reverse complement.

When the libraries are cloned, the crRNA cassette consists of 4 multiplexed crRNAs, but the oligo nucleotides were designed to only include 2 crRNAs and combine during cloning with another oligonucleotide to form the complete 4 crRNA expression cassette. This allows to order shorter-sized oligonucleotides as well as improve flexibility of combinations and ultimately greatly reduce pool size since it is not necessary to incorporate pairwise combinations within it. Each targeted gene has two oligonucleotide versions, the front-half and the tail-half versions in this disposition:

5’-[Forward primer]**CGTCTC**AAGAT(DRwt)[crRNA1.1](DR1)[crRNA1.2]GGAAA**GAGACG**[Reverse primer]-3’
5’-[Forward primer]**CGTCTC**AGGAA(DR3)[crRNA2.1](DR6)[crRNA2.2]TTTTTTGAATC**GAGACG**[Reverse primer]-3’

In bold, the recognition sequence of the endonuclease type IIS BsmBI 5’-CGTCTC-3’ or 3’-GCAGAG-5’. Underlined are the overhangs generated by BsmBI digestion. DR stands for Direct Repeat, several DR versions were used to reduce the chances of recombination events in the lentiviral production step (adopted from DeWeirdt et al ^39^). A six consecutive thymine stretch RNA Polymerase III termination signal was included at the end of the tail-half oligonucleotides. Specific primers flank the oligonucleotides to obtain double stranded DNA and allow for separate amplification of the different sets of oligonucleotides included in the pool.

The oligonucleotide pool was procured from Twist Biosciences as single-stranded DNA. Full-length oligonucleotide sequences, crRNAs with Direct Repeats and primers used in this article are found in **[Table S2]**.

### Modules extraction and cloning

The MMR, BER and DR targeting modules were amplified from the oligonucleotide pool in separate PCR reactions and run in triplicate. Each PCR was performed using KAPA HiFi 2X HotStart ReadyMix (Roche) using 1 ng of starting template (oligo pool) per 25 μl reaction using the module-specific primer pairs at a final concentration of 0.3 μM. The thermocycler program was as follows: denaturation at 95°C for 3 min, followed by 12 cycles of 20 s at 98°C, 30 s at 62°C, and 30 s at 72°C, followed by a final extension of 1 min at 72°C. Triplicates were then merged for the next steps. The PCR products and the backbone (Addgene, pRDA_052) were digested with BsmBI_v2 (NEB, #R0580), followed by gel purification using NucleoSpin Gel and PCR clean-up, MACHEREY-NAGEL.

Golden Gate Assembly was performed in triplicates using BsmBI_v2 and T4 DNA ligase (NEB, #M0202) in a one-pot reaction. Specifically, the assembly reaction consisted of 330ng linearized vector, 50 ng of each library module PCR, 3 μl of NEBridge Golden Gate Assembly kit (NEB, #E1602), 3 μl of T4 DNA ligase buffer 10X, and nuclease-free water to a final volume of 30 μl. The reaction was cycled between 5 min at 37°C (digestion) and 5 min at 16°C (ligation) for 35 cycles, with a final incubation at 60°C for 10 minutes.

Following assembly, the reaction mixture was transformed into chemically competent OmniMAX cells (ThermoFisher, C854003) via heat shock at 42°C for 30 seconds. Transformed cells were plated onto 10 LB agar plates containing 100 μg/mL of ampicillin and incubated overnight at 33°C. The final plasmid library was purified using a maxiprep kit (QIAGEN Plasmid Maxi Kit, Qiagen). Sequencing of the final libraries showed high cloning fidelity, with 94% of reads matching expected combinations in the MMR-BER/direct-repair library used for this study (other, unrelated gene sets were included in the same oligonucleotide pool).

### Transduction

Cell lines stably expressing enCas12a or the crRNA library were obtained by lentiviral transduction. 3.8×10^6^ HEK293T cells were seeded in a 10 cm dish the day prior to transfection to achieve 80% confluency. Transfection was performed using PEI following the manufacturer’s instructions, with a plasmid ratio of 4:3:1 for the library vector, psPAX2 (packaging plasmid), and pMD2.G (envelope plasmid). The transfection mixture was incubated with cells for 6 hours followed by replacement with DMEM supplemented with 10% FBS. Supernatants containing lentiviral particles were collected at 48 hours post-transfection and filtered through a 0.45 μm syringe filter. Viral titers were estimated by infectivity assay on the target cells (K562 cells).

48h post-enCas12a transduction, K562 cells were selected with blasticidin (at 12 µg/ml final concentration) during 14 days. Cells were then single-cell sorted and after expansion, the clones were checked for enCas12a expression by Western blot using H-tag antibody.

48h post-crRNA library transduction, the bulk-transduced K562 cell population was selected with puromycin (at 1 µg/ml final concentration) during the next 72h. Cells were then single-cell sorted and after expansion, the clones were checked for crRNA cassette incorporation by amplicon sequencing.

### DNA extraction

For checking integrated crRNAs and CRISPR edits, genomic DNA was extracted from edited and control samples using PrepGEM Universal kit (Microgem). Specifically, roughly 10^4^ cells were pelleted and resuspended in 2.5 μl of 10x Buffer BLUE, 0.5 μl of prepGEM and 27 μl of distilled nuclease-free water. The mix was vortexed and then incubated for 5 min at 75°C and 2 min at 95°C before use.

For WGS, genomic DNA was extracted from the K562 isogenic clones using the Mag-Bind Blood & Tissue DNA HDQ 96 kit (OMEGA Bio-tek), following the manufacturer’s protocol. Briefly, 10^6^ cells were harvested and washed with sterile phosphate-buffered saline (PBS). Cell pellets were lysed in 180 μl of PBS with proteinase K and incubated at 55°C for 10 minutes. DNA was bound to silica-based beads washed and purified on a magnet and eluted in 50 μl of nuclease-free water.

### Checking crRNA combination and enCas12a edits

The crRNA cassette was amplified by PCR using flanking primers that anneal to the plasmid backbone. The genotyping PCRs were performed using Biotools Master Mix 2X (Biotools) using 0.5 μl of the PrepGEM lysate in 7 μl reactions with the Argon and Beaker primers (shown below) at a final concentration of 0.3 μM. The thermocycler program was as follows: denaturation at 95°C for 5 min, followed by 35 cycles of 20 s at 98°C, 30 s at 56.5°C, and 30 s at 72°C, followed by a final extension of 3 min at 72°C.

Argon FW (5’-TTGTGGAAAGGACGAAACACCG-3’)
Beaker RV (5’-CCAATTCCCACTCCTTTCAAGACCT-3’)

The resulting crRNA cassette amplifications were Sanger sequenced (Macrogen) using the same “Beaker RV” primer described above for the sequencing. The target genomic region flanking the edited site was amplified by PCR using primers designed to encompass the regions of interest (primer sequences provided in **[Table S3]**). The PCR reactions were performed using Biotools Master Mix 2X (Biotools) using 0.5 μl of the PrepGEM lysate in 7 μl reactions with their respective primers at a final concentration of 0.3 μM. The thermocycler program was as follows: denaturation at 95°C for 5 min, followed by 35 cycles of 20 s at 98°C, 30 s at 60°C, and 30 s at 72°C, followed by a final extension of 3 min at 72°C.

Purified PCR products were barcoded using Oxford Nanopore Technologies’ Native Barcoding kit 96 (SQK-NBD110.96 and SQK-NBD114.94) following the manufacturer’s protocol and the sequencing libraries were prepared using Oxford Nanopore Technologies’ Ligation Sequencing Kit (SQK-LSK110 and SQK-LSK114). Libraries were loaded onto Flongle cells (Oxford Nanopore Technologies, FLO-FLG001 and FLO-FLG114) for sequencing.

Sequencing reads were aligned using Minimap2 ^77^ to the expected WT amplicons taking the GRCh38 genome as reference. Alignment data was visualized using IGV ^78^ to assess whether all alleles within the amplified region were fully edited.

### DNA sequencing, alignment and variant calling

Library preparation and DNA sequencing was done on Illumina NovaSeq 6000 machines at the Sequencing Unit of Centro Nacional de Análisis Genómico (CNAG-CRG), Barcelona, Spain or at Genomic Medicine, Rigshospitalet, Copenhagen, Denmark, using 2×150 paired end reads, with an average sequencing depth of 28.3x (minimal WGS coverage of 14.5x for a MUTYH-NEIL2 clone, the remainder in the range of 18.4-67x). Illumina reads were aligned to the GRCh38.d1.vd1 reference genome using the BWA-MEM algorithm (v0.7.17). Germline variant calling was performed using Strelka2 (v2.9.10) in multi-sample mode to identify substitutions and InDels. The resulting multi-sample VCF was annotated using ANNOVAR (version 2019Oct24) with the gnomAD hg38 genome database, and variants with a gnomAD_genome_ALL allele frequency ≥ 0.001 were excluded to remove common polymorphisms from the callset. Variants outside uniquely mappable regions, as defined by the k50 Umap mappability track, were additionally removed to minimise the inclusion of alignment artefacts. To define a set of clone-discrete variants for each genotype, the multi-sample VCF was further filtered to retain only variants genotyped in no more than 3 clones. Variants were required to pass Strelka2 quality filters, with empirical variant score (EVS) thresholds of ≥ 7 for SNVs and ≥ 6 for InDels, a QUAL score of ≥ 5, and a minimum of 3 supporting reads per clone. To compare mutation distributions to genomic features, we used the *wgEncodeUwRepliSeqK562WaveSignalRep1_GRCh38.bed* replication timing dataset and the ENCFF649RBS whole genome bisulfite sequencing dataset generated from CD34+ myeloid progenitor tissue, both from Encode.

### Short tandem repeat calling and spectrum generation

Short tandem repeat indels were called using repeat allele length histograms generated by MSISensor ^79^, a specialized tool for accurate STR indel detection. MSISensor was run using default settings, using the wild-type K562 sample as the reference. The histograms were analysed using a bespoke calling method. Briefly, for each locus, simulated homo- and heterozygous histograms were generated for 1, 2 or 3 units long deletions and insertions by shifting the peaks of the observed normal histogram accordingly. The best fit of the observed sample histogram was selected by a maximum likelihood estimation approach, and empirical p-values were calculated by bootstrapping on the observed normal histogram peaks. For the spectrum generation, only uniquely mappable loci were used according to the GRC Alignability 50 track, and only indels with a log-likelihood ratio difference of min. 10 were retained comparing the normal histogram vs. the optimal model. For the spectrum generation, several STR-relevant features were used for categorization: repetition length capped at 25x, repeat unit length and sequence, indel unit length and type (deletion/insertion), binarized replication timing and overlap with Alu elements.

### Non-negative matrix factorization (NMF)

NMF was utilized to find latent components in the short nucleotide variation (SNV) and indel mutations of the K562 cell line panel. The SNVs and indels were subjected to NMF in a joint manner, i.e., the COSMIC SBS96-formatted SNV and the COSMIC ID83-formatted indel spectra were concatenated for each sample. To extract signatures, three different NMF decomposition tools were used, SigProfiler ^80^, MuSiCal ^81^ and MUSE-XAE ^82^. NMF rank values between 2 and 20 were tested with each tool, and for each rank, silhouette stability and root mean squared difference (RMSD) values were collected. While the automatic rank suggestions of the three tools were k=6, 7 and 6, respectively, we proceeded for k=11 of the SigProfiler solution in order to detect low intensity components or ones only present in a low number of samples. Refitting of the isolated components on the batch effect corrected RPE1 and iPSC cell line samples was performed using MuSiCal, with the “likelihood_bidirectional” method and a threshold of 0.001.

### Elastic net-based MMR classification

We used an Elastic net based modeling framework to construct a model capable of correctly predicting MMR genotypes based on the joint SNV+indel SBS179 spectra, with the K562 cell line panel taken as the training set. The model estimated the probability of all seven possible MMR labels (*MSH2*, *MSH6*, *MLH1*, *PMS2*, *MSH3*, *MLH3* and MMR_proficient) concurrently. The Elastic net was run with parameters family = “multinomial”, alpha = 0.5 and type_multinomial = “grouped”, in a 100x bootstrap. The predictor variables also included the total SNV burden, the indel burden, the SNV/indel ratio and the estimated age of the K562 samples. This pseudo-age was needed to control for the highly different mutation accumulation times between the cell line based mutation spectra and the cancer cases used later, and was estimated by a negative binomial regression using the glm R package, based on the known ages of the TCGA patients. Before running the predictions, the predictor variables were scaled, and 100-fold bootstrapping was used to obtain Bias-Corrected and Accelerated confidence intervals.

We devised two measures based on Z-score calculations to express the uncertainty of model predictions. Z_def_ was defined as the difference of the probabilities of the top and the second best label prediction, divided by the bootstrap confidence interval of the top label, and was used to assess the general confidence of the genotype-level final prediction. Z_prof_ was defined as the difference between the top predicted and the “MMR_proficient” label, divided by the confidence interval of the latter, and described the confidence of the model about the given sample being MMR proficient or not.

### MMR genotype-specific STR locus selection

The initial STR set containing 1.795.190 loci was first limited by removing non-uniquely mappable sites and all loci that were mutated in less than three K562 samples, generating a basic candidate list of 815243 sites. Next, to reduce overfitting, an Elastic net-based pre-filtering was applied. Briefly, each targeted genotype (*MSH2*, *MSH3*, *MSH6*, *MLH1* and *PMS2*) was treated as a binary variable, and regularized logistic regressions were used with an alpha parameter of 0.05 (biased towards ridge regression, thus retaining more potential sites). To control stochasticity and overfitting, the lambda parameter was determined by 100 instances of 5-fold cross-validation, and the 10th percentile of the lambda distribution was used. The final Elastic net step consisted of a leave-one-out cross-validation (LOOCV) performed across the whole K562 panel (n = 71) in order to select robust, stable loci, and only STR sites found in at least 90% of the LOOCV iterations were retained.

Next, to prioritize among the several hundreds of loci per genotype, we employed an independent odds ratio-based ranking method. Briefly, the genotypes were again treated as binary variables, and for each candidate STR locus, odds ratios were calculated based on 2×2 contingency tables (locus mutated/non-mutated vs sample belongs/not to target genotype), with the Haldane-Anscombe correction, separately for the K562 and the Lynch patient cohort. Finally, the minimum of the odds ratio was taken, and for each genotype, the number of loci sufficient to achieve 90% sensitivity was selected as the final MMR-genotype-specific locus set.

### External cell line datasets

For external validation, we utilized two external datasets containing mutational spectra of isogenic DNA repair mutant panels: RPE1 cells from Koh et al ^53^ and induced pluripotent stem cells (iPSCs) from Zou et al ^2^. Both cell line panels were filtered for genotypes overlapping with our K562 set [**Table S8**]. To harmonize differences due to different cell types, maintenance conditions etc., we performed batch effect correction using ComBat from the R package sva-3.56.0 ^83^ using the known genotypes as additional covariates.

### TCGA cases

We tested our MMR genotype prediction method on TCGA projects, TCGA-COAD, -READ, - UCEC and -STAD. In order to enrich MSI-H and MSI-L samples in the final validation cohort [**Table S9**], we used the Bethesda MSI panel-based PCR assessments from the tissue-wise nationwidechildrens.org_auxiliary_*.txt files, available as parts of TCGA metadata. Variant calling with Strelka and MSISensor and posterior filtering in the downloaded normal and tumor datasets was performed identically as in the K562 cell line panel. A two-step batch effect correction pipeline was used (applied on the square-root transformed mutation counts) to harmonize the spectra from the WGS datasets firstly among TCGA cases from different tissues, and next between spectra in WGS of K562 cells and TCGA cases.

In order to remove out-of-distribution samples from the TCGA cohort, we used the mclust R package ^84^ to cluster the TCGA spectra, and selected only clusters showing a high similarity to MMR proficient and MMR deficient in the K562 panel. The samples filtered here (mostly polymerase point mutant and polymerase/MMR double mutant cases) were used for a separate, out-of-distribution validation analysis of the MMRd Elastic net models. To prevent these out-of-distribution, hypermutated samples distort the batch effect correction pipeline necessary for applying Elastic net predictions to the TCGA data, the batch effect correction and scaling were performed using parameters taken from the batch effect correction step of *bona fide* MMR proficient TCGA samples.

Lynch syndrome patients were identified by performing variant calling on the paired normal-tumor pairs with Strelka, and selecting coding variants in MMR genes (MSH2, MLH1, PMS2, MSH6, MSH3) that a) were frameshift indels, stop-gain mutations or missense variants with AlphaMissense scores higher than 0.6 and b) were clonally present in both the normal and tumor sample WGS of a given TCGA patient, implying they were germline variants.

## Supplemental information

**PDF Document. Supplementary Figures S1-S18.**

**Spreadsheet. Supplementary Table S1-S10.**

