## Supplementary Figure S1-S18 for "Multidimensional mutational phenotypes of MMR deficiency in human cancer cells"

### McCullough, Poti et al (2026) Supplementary Figures S1-S18

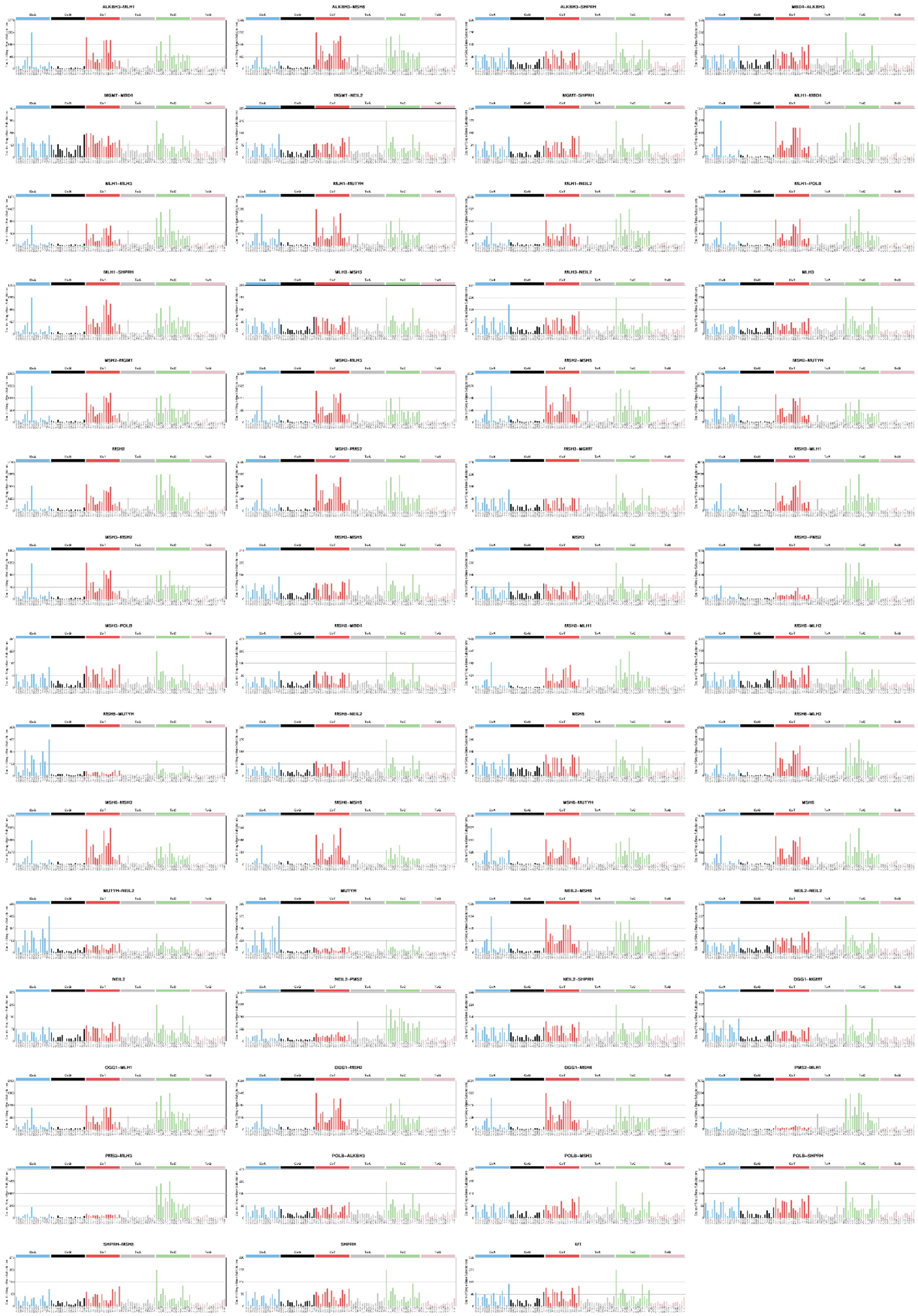

**Supplementary figure S1. SNV (SBS) mutational count spectra of the K562 cell line panel.**

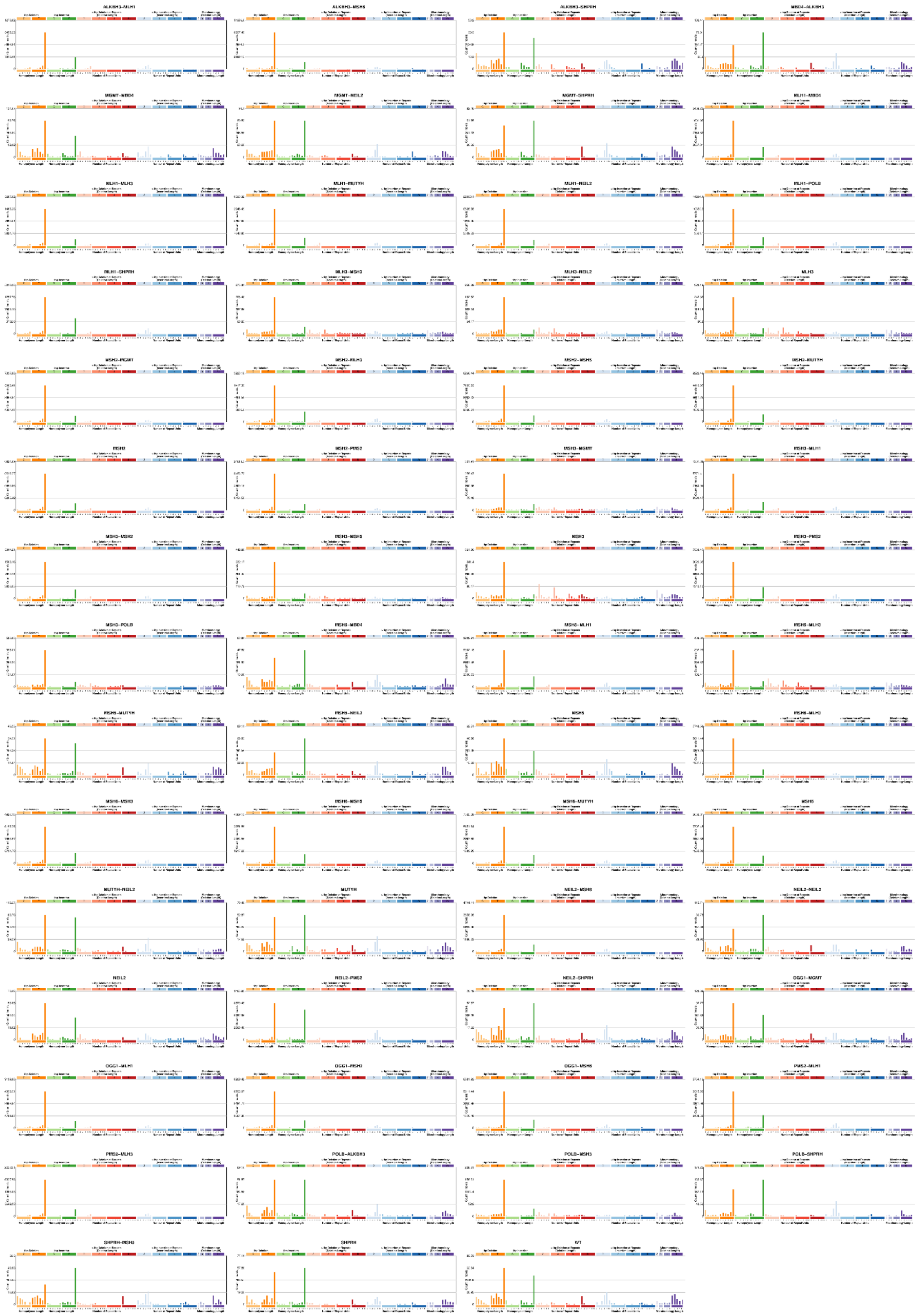

**Supplementary figure S2. Indel mutational count spectra of the K562 cell line panel.**

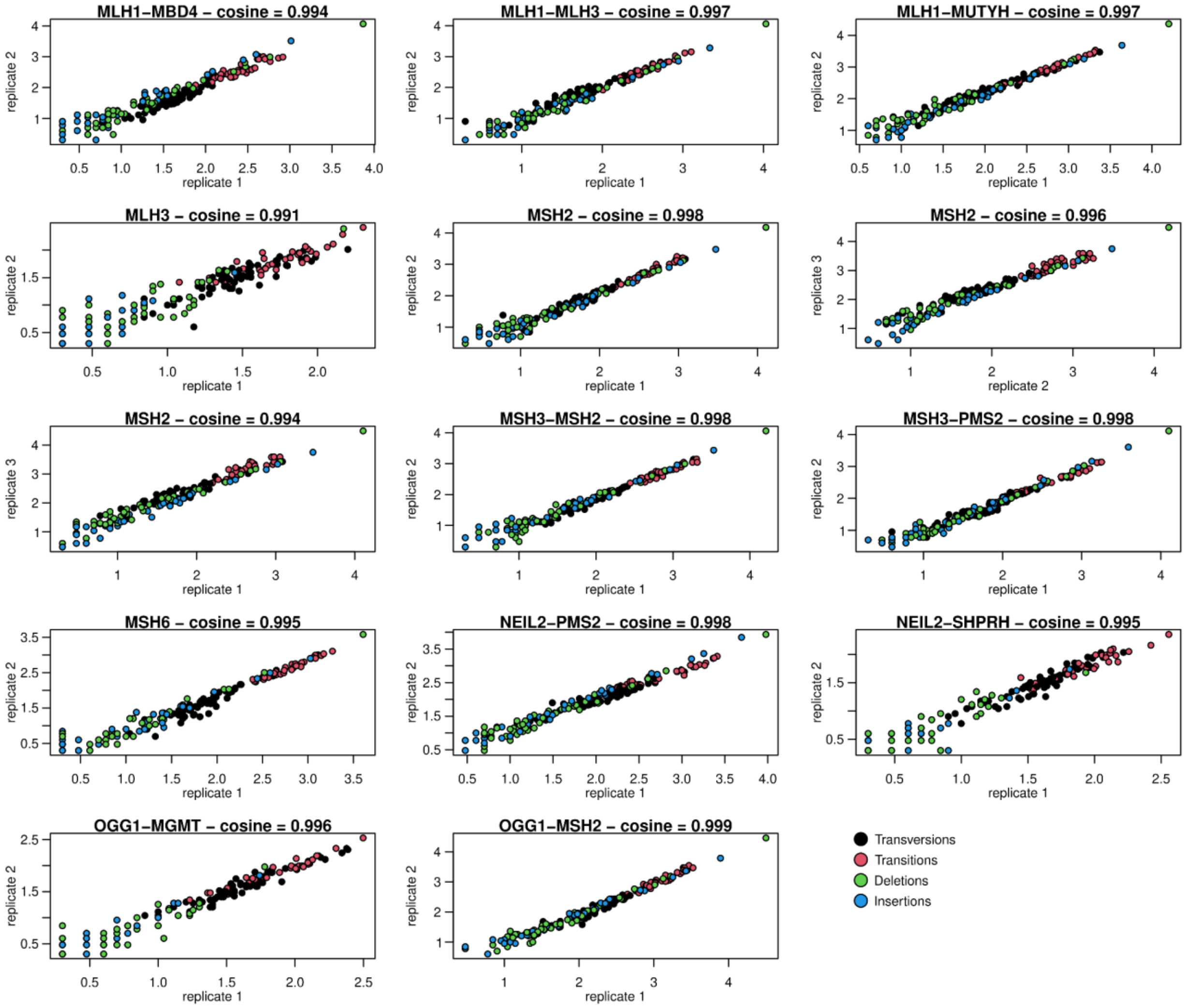


**Supplementary figure S3. Reproducibility of mutational spectra across biological replicates of gene ablation and mutation accumulation.** Correspondences of mutational counts in  the COSMIC SBS96+ID83 categories of genotypes with more than one replicate clones. Cosine similarities are annotated.

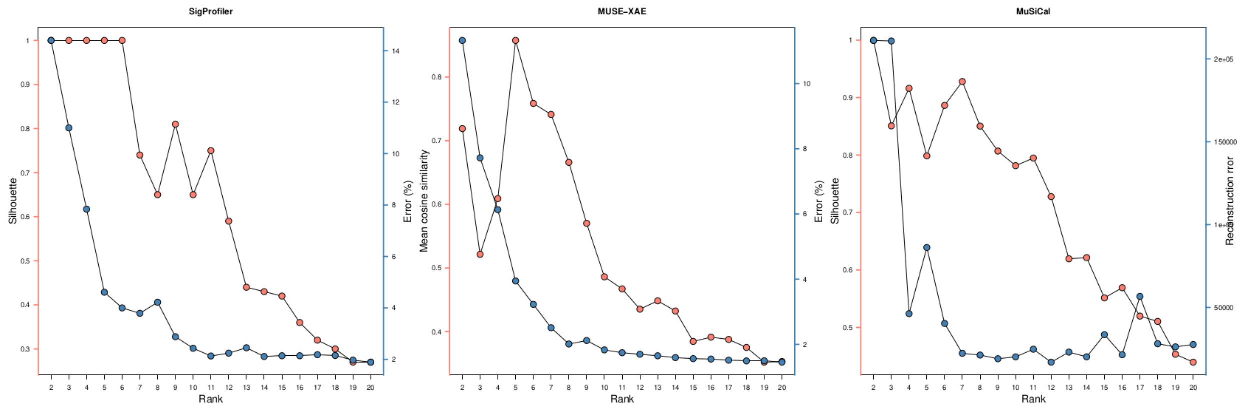


**Supplementary figure S4. NMF rank estimation of the K562 WGS dataset, using silhouette index stability and reconstruction error (RMSD) as readouts, for different mutational signature tools.**

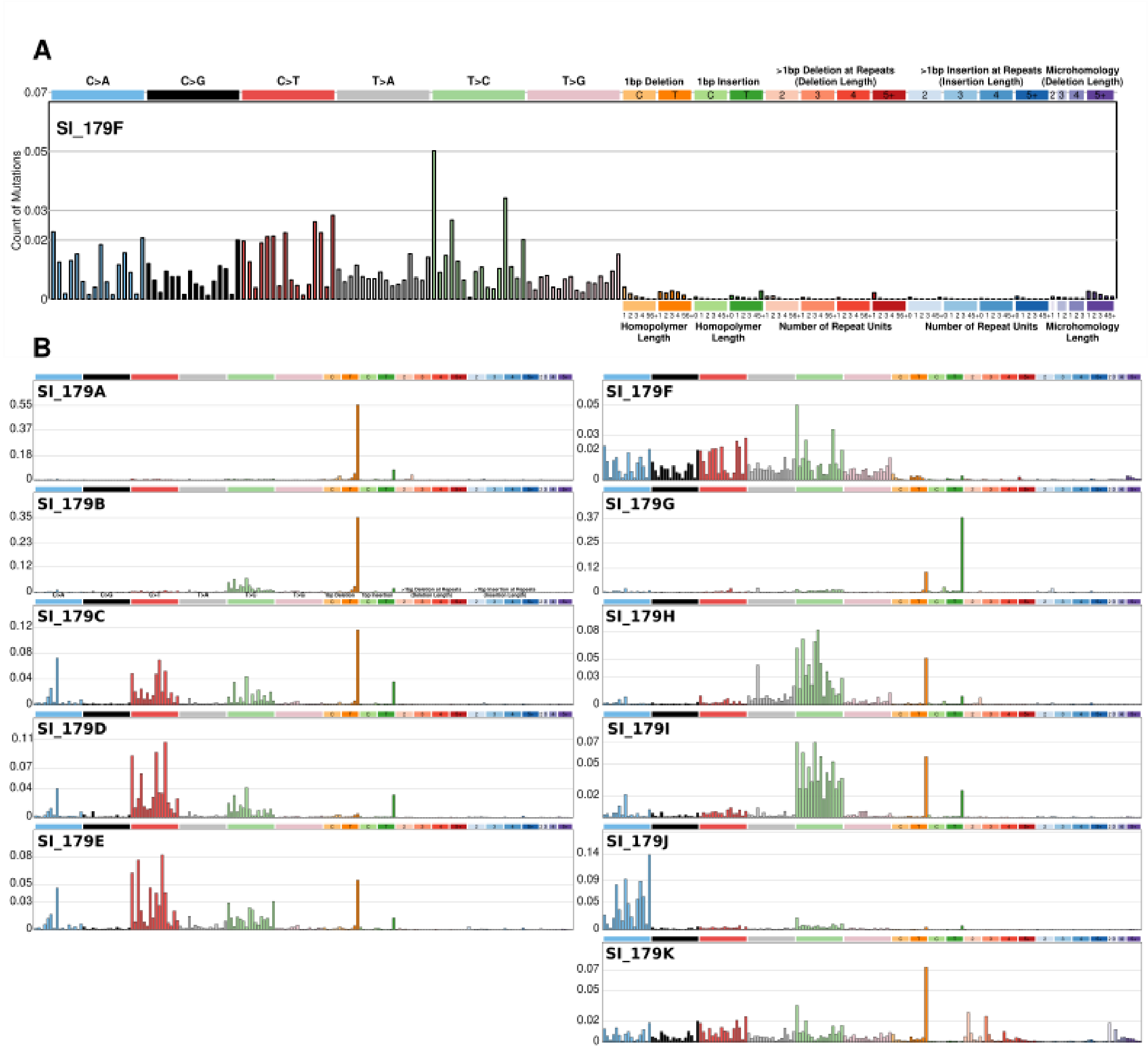


**Supplementary figure S5. NMF signatures identified from the K562 cell line panel of the present study. A.** The structure of the extracted NMF components: COSMIC-style SNV and indel categories were concatenated. SI_179F was identified as the background mutational pattern in K562 cells, with its SBS spectrum resembling COSMIC SBS5 most strongly, and additionally SBS40a/c and SBS92 (see Supplementary Fig. S7). **B.** The spectra of the 11 extracted NMF signatures.

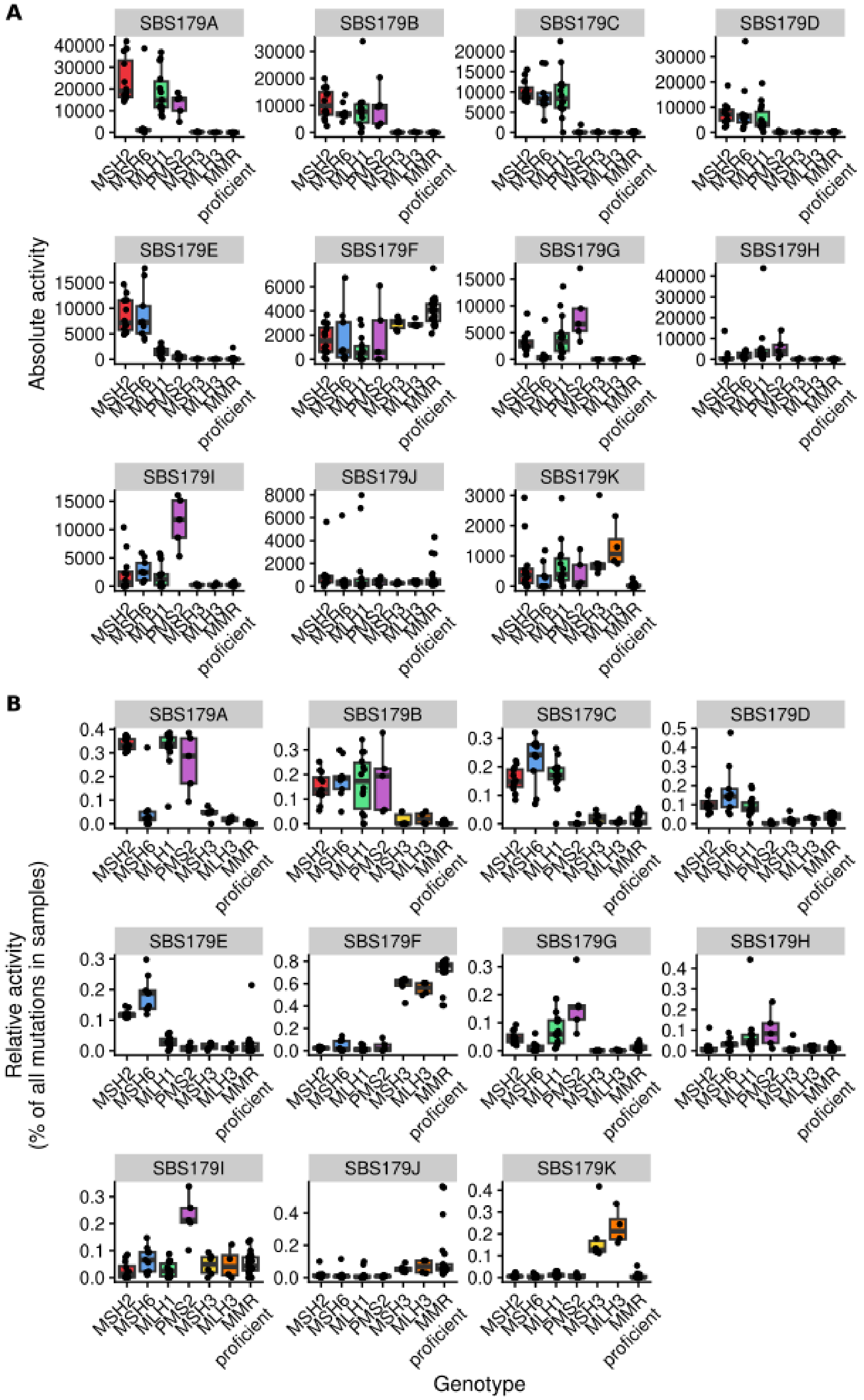


**Supplementary figure S6. NMF signature activities in the K562 cell line panel. A.** Absolute activities, the number of mutations attributed to each signature in each sample. **B.** Relative activities, highlighting the relative influence of each signature in each sample.

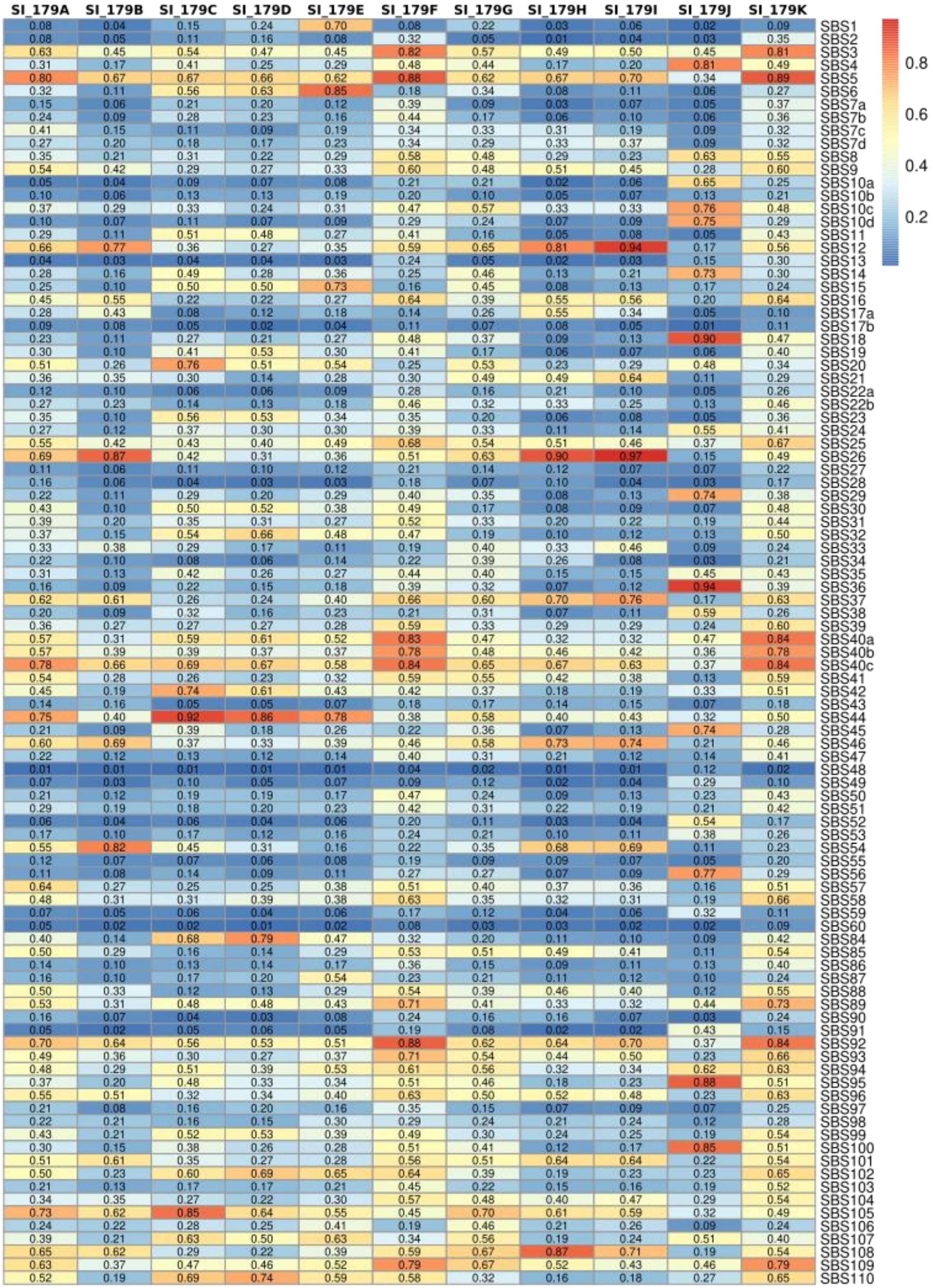


**Supplementary figure S7. Cosine similarities of the SNV components of NMF signatures identified in this study against the COSMIC SBS v3.5 signature set.**

**A**

| | SI_179A | SI_179B | SI_179C | SI_179D | SI_179E | SI_179F | SI_179G | SI_179H | SI_179I | SI_179J | SI_179K |
|---|---|---|---|---|---|---|---|---|---|---|---|
| ID1 | 0.14 | 0.05 | 0.29 | 0.97 | 0.23 | 0.31 | 0.96 | 0.18 | 0.40 | 0.81 | 0.01 |
| ID2 | 0.99 | 1.00 | 0.96 | 0.16 | 0.97 | 0.03 | 0.27 | 0.97 | 0.92 | 0.52 | 0.83 |
| ID3 | 0.06 | 0.04 | 0.07 | 0.16 | 0.09 | 0.51 | 0.10 | 0.07 | 0.07 | 0.23 | 0.12 |
| ID4 | 0.01 | 0.00 | 0.01 | 0.01 | 0.03 | 0.19 | 0.01 | 0.03 | 0.01 | 0.09 | 0.51 |
| ID5 | 0.18 | 0.17 | 0.15 | 0.10 | 0.19 | 0.70 | 0.06 | 0.19 | 0.15 | 0.24 | 0.30 |
| ID6 | 0.03 | 0.03 | 0.04 | 0.06 | 0.05 | 0.48 | 0.05 | 0.04 | 0.04 | 0.17 | 0.10 |
| ID7 | 0.11 | 0.06 | 0.07 | 0.21 | 0.10 | 0.13 | 0.06 | 0.10 | 0.04 | 0.06 | 0.08 |
| ID8 | 0.04 | 0.03 | 0.05 | 0.08 | 0.05 | 0.59 | 0.06 | 0.04 | 0.06 | 0.16 | 0.13 |
| ID9 | 0.04 | 0.04 | 0.03 | 0.07 | 0.08 | 0.71 | 0.03 | 0.04 | 0.03 | 0.15 | 0.14 |
| ID10 | 0.01 | 0.00 | 0.01 | 0.03 | 0.03 | 0.19 | 0.01 | 0.02 | 0.01 | 0.06 | 0.10 |
| ID11 | 0.03 | 0.02 | 0.04 | 0.08 | 0.04 | 0.11 | 0.08 | 0.05 | 0.05 | 0.09 | 0.03 |
| ID12 | 0.08 | 0.03 | 0.02 | 0.05 | 0.04 | 0.21 | 0.03 | 0.15 | 0.02 | 0.05 | 0.12 |
| ID13 | 0.03 | 0.02 | 0.03 | 0.05 | 0.05 | 0.42 | 0.04 | 0.03 | 0.03 | 0.16 | 0.08 |
| ID14 | 0.14 | 0.11 | 0.21 | 0.48 | 0.23 | 0.27 | 0.47 | 0.18 | 0.28 | 0.43 | 0.09 |
| ID15 | 0.19 | 0.18 | 0.17 | 0.13 | 0.19 | 0.49 | 0.05 | 0.17 | 0.15 | 0.20 | 0.20 |
| ID16 | 0.02 | 0.01 | 0.05 | 0.15 | 0.06 | 0.15 | 0.15 | 0.06 | 0.07 | 0.15 | 0.03 |
| ID17 | 0.01 | 0.00 | 0.01 | 0.04 | 0.03 | 0.16 | 0.03 | 0.02 | 0.02 | 0.07 | 0.05 |
| ID18 | 0.24 | 0.24 | 0.22 | 0.10 | 0.25 | 0.59 | 0.08 | 0.23 | 0.20 | 0.22 | 0.30 |
| ID19 | 0.03 | 0.02 | 0.03 | 0.10 | 0.08 | 0.27 | 0.06 | 0.07 | 0.04 | 0.11 | 0.07 |
| ID20 | 0.04 | 0.03 | 0.06 | 0.13 | 0.06 | 0.06 | 0.15 | 0.09 | 0.07 | 0.11 | 0.03 |
| ID21 | 0.15 | 0.14 | 0.15 | 0.10 | 0.17 | 0.22 | 0.11 | 0.16 | 0.16 | 0.19 | 0.53 |
| ID22 | 0.03 | 0.03 | 0.04 | 0.06 | 0.05 | 0.07 | 0.06 | 0.04 | 0.05 | 0.08 | 0.04 |
| ID23 | 0.04 | 0.03 | 0.03 | 0.06 | 0.07 | 0.65 | 0.02 | 0.05 | 0.04 | 0.18 | 0.22 |
| ID24 | 0.03 | 0.02 | 0.03 | 0.07 | 0.05 | 0.27 | 0.05 | 0.05 | 0.03 | 0.15 | 0.30 |
| ID25 | 0.22 | 0.15 | 0.33 | 0.79 | 0.31 | 0.50 | 0.79 | 0.26 | 0.42 | 0.75 | 0.17 |

Color scale: 0, 0.2, 0.4, 0.6, 0.8, 1

**B**

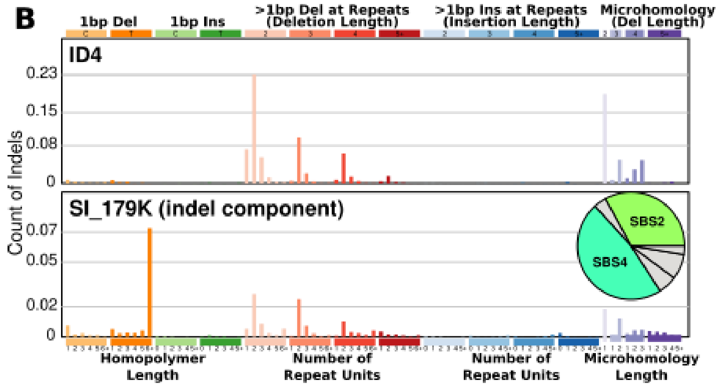


**Supplementary figure S8. A.** Cosine similarities of the indel components of NMF signatures identified in this study against the COSMIC ID v3.5 signature set. **B.** Direct comparison of the indel component of SI_179K and COSMIC ID4. The inlay shows the estimated contributions of COSMIC ID signatures in SI_179.

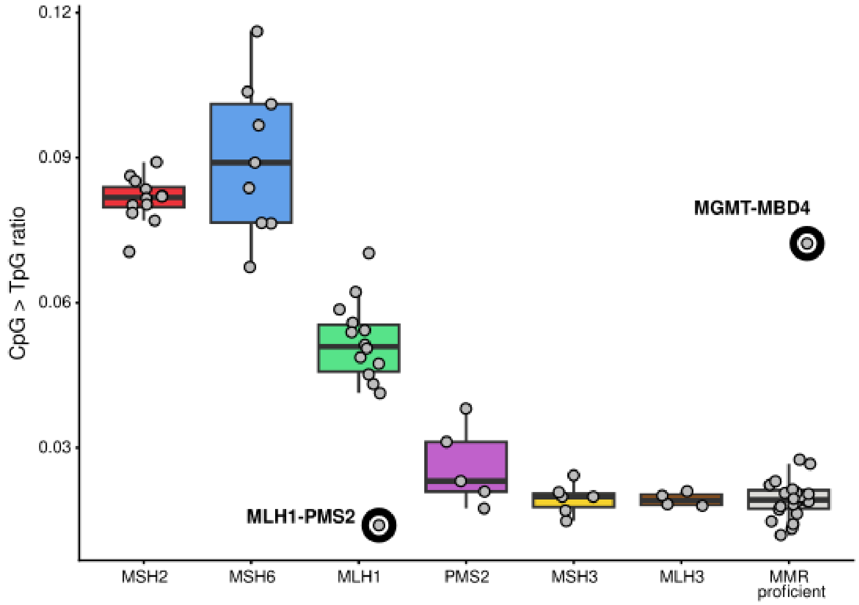


**Supplementary figure S9. Ratio of CpG > TpG SNV counts and total SNV burden**, grouped by MMR genotypes present in the study. Two outlier samples due to specific genotype interactions are marked by thick circles.

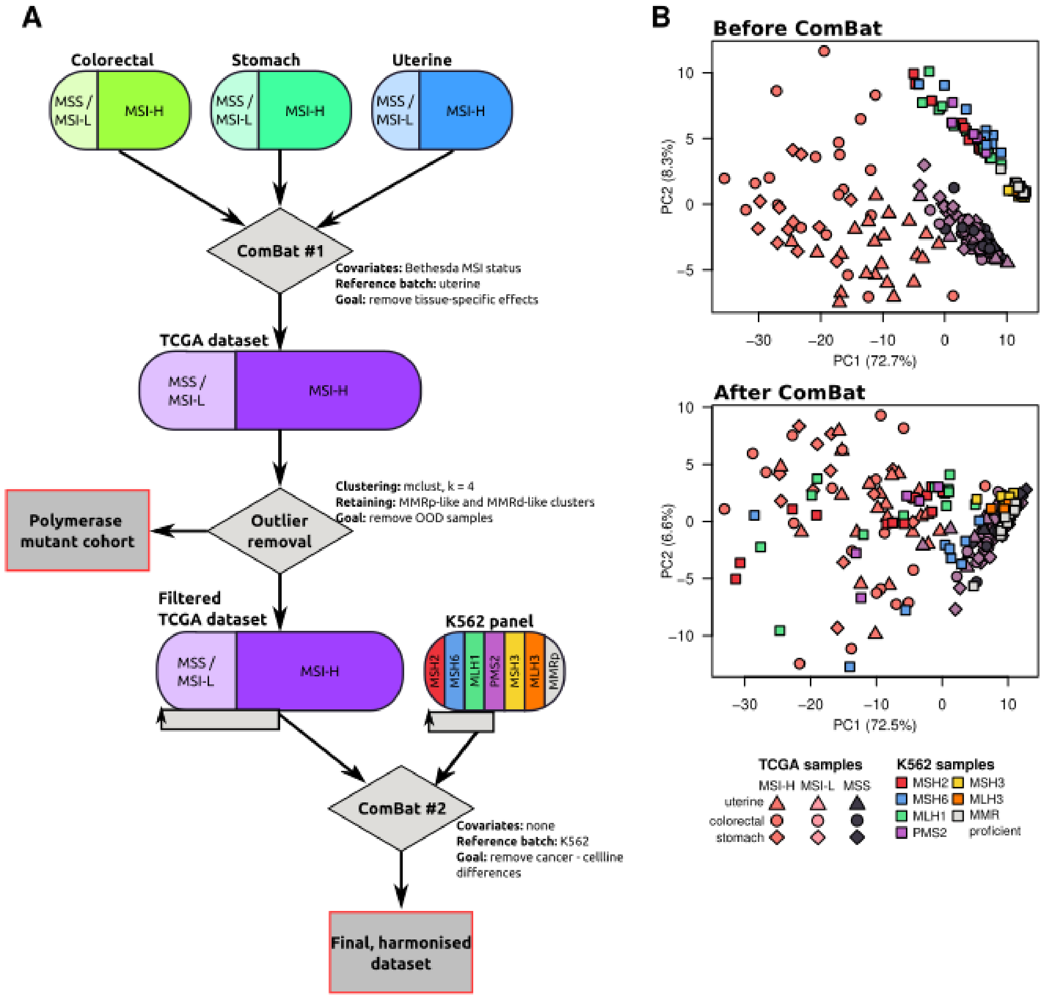


**Supplementary figure S10. Batch effect correction between the K562 panel and the MSI-enriched TCGA dataset. A.** Overview of the harmonisation pipeline for the TCGA dataset. **B.** First two principal components of the joint SBS+ID spectra of K562 and TCGA samples, before and after batch effect correction, showing an improved overlap between the two groups.

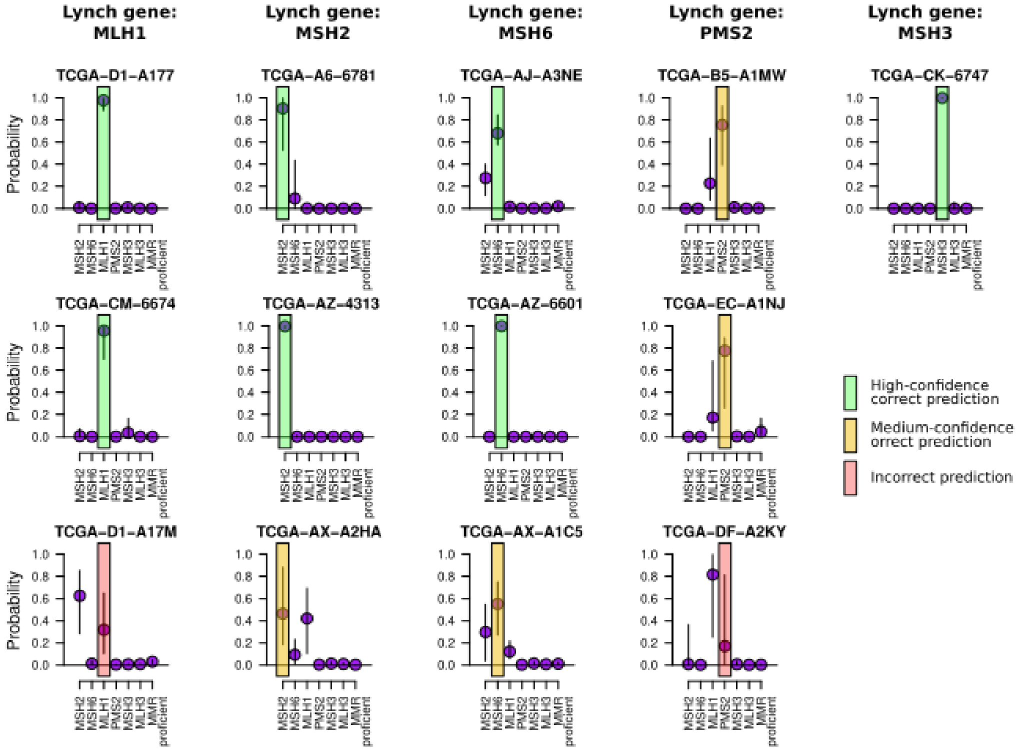


**Supplementary figure S11. Detailed, per-genotype Elastic net model predictions for the 13 Lynch patients identified in the TCGA data selection.** The colored shading indicates if the model prediction was incorrect, and if the model confidence assessed by the $Z_{def}$ score was high- or medium-level.

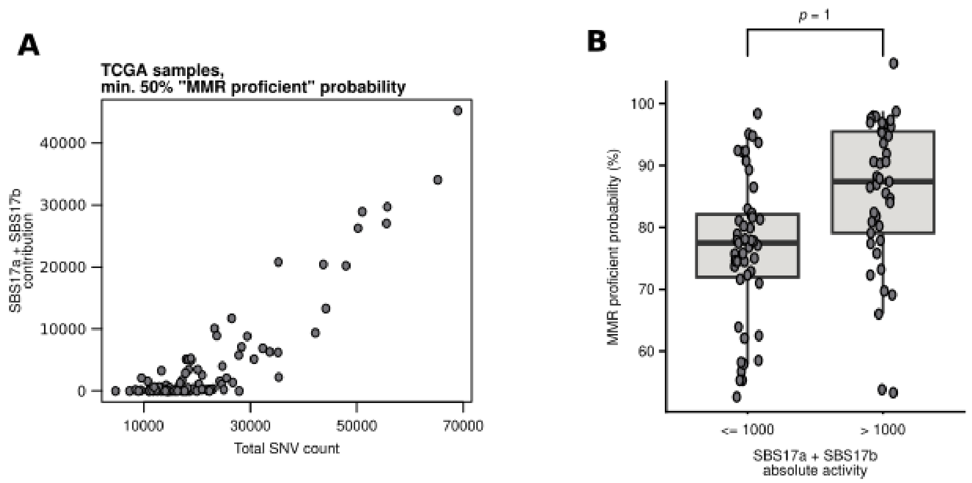


**Supplementary figure S12. Effect of SBS17a and SBS17b hypermutation on the Elastic net prediction of MMRd. A.** Association between SBS17 activity and total SNV counts in samples with at least 50% probabilities of MMR proficiency, showing that SBS17 is the dominant mutagenic effect in MMR proficient cases in the TCGA cohort. **B.** Difference in prediction probabilities of the “MMR proficient” label in cases with or without min. 1000 SBS17 associated SNVs (one-tailed Wilcoxon-test, H0: ‘> 1000’ less than ‘<= 1000‘)

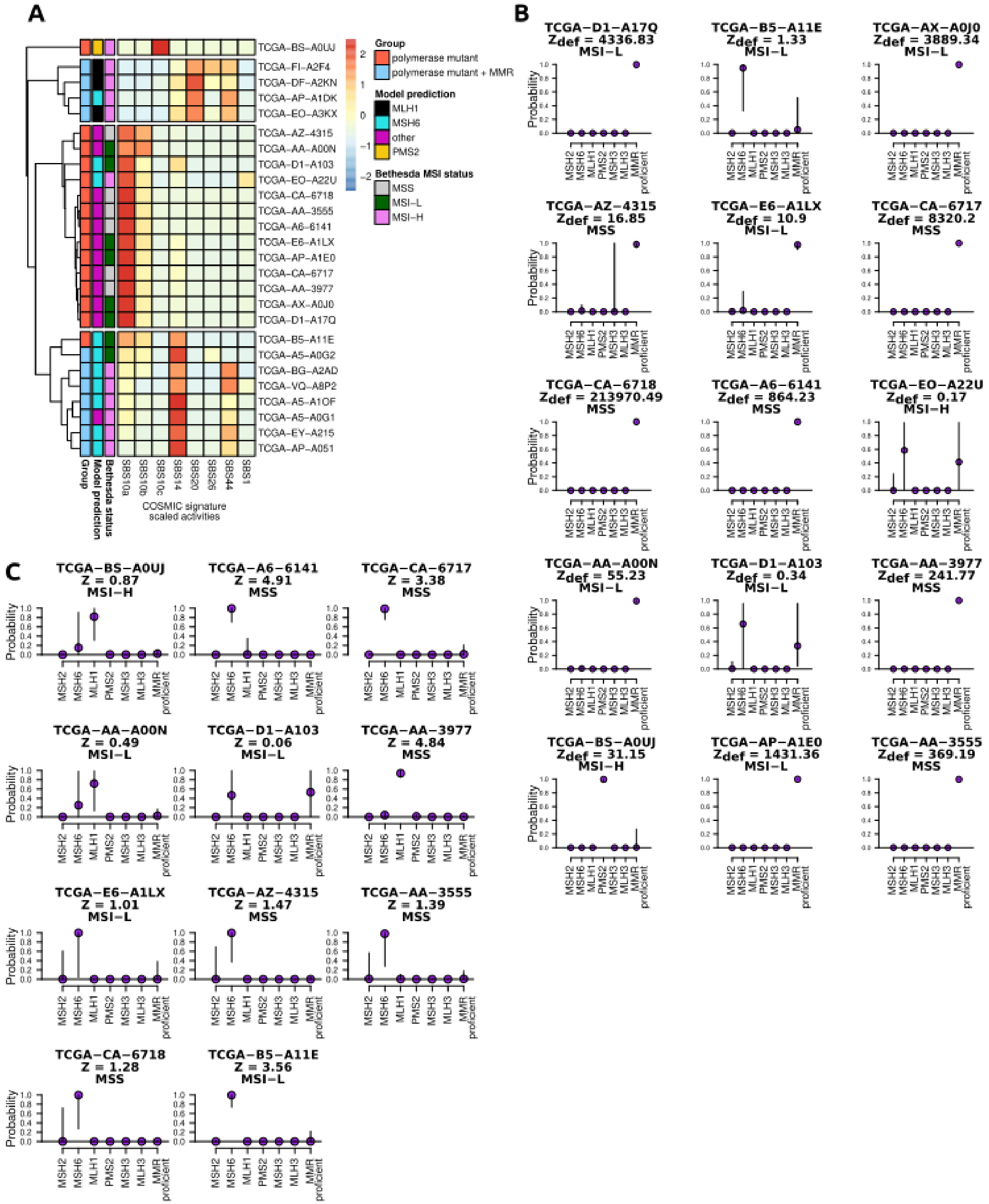


**Supplementary figure S13. Assessment of Elastic Net models' performance on out-of-distribution WGS, with hypermutated but presumably not MMRd samples. A.** Comparison of model performances, Bethesda MSI annotations and COSMIC SBS96 signature activities in the identified DNA polymerase mutant or DNA polymerase+MMR mutant cases in the TCGA cohort. **B-C.** Detailed model predictions +/- confidence intervals for DNA polymerase mutant (B) and DNA polymerase + MMR mutant TCGA cases (C).

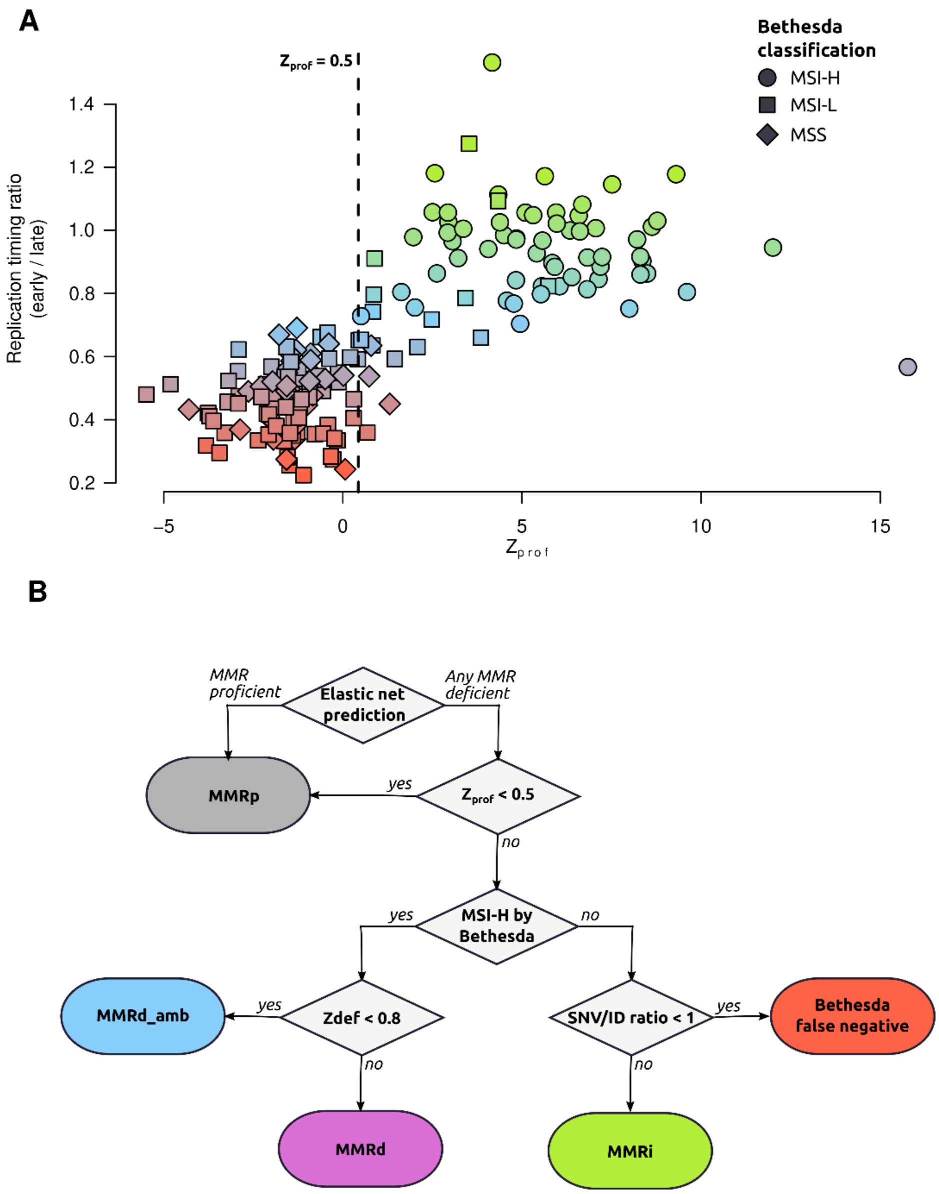


**Supplementary figure S14. Classification of MMR intermediate (MMRi) samples. A.** Relationship between Zprof scores and replication timing ratios. The marker colors refer to bins of replication timing ratios, and marker shapes refer to Bethesda PCR classifications of each TCGA sample. The dashed line at Zprof = 0 indicates the threshold used for separating MMRi vs MMRd cases. **B.** Re-classification algorithm of the cancer cases in MSI-enriched TCGA subset.

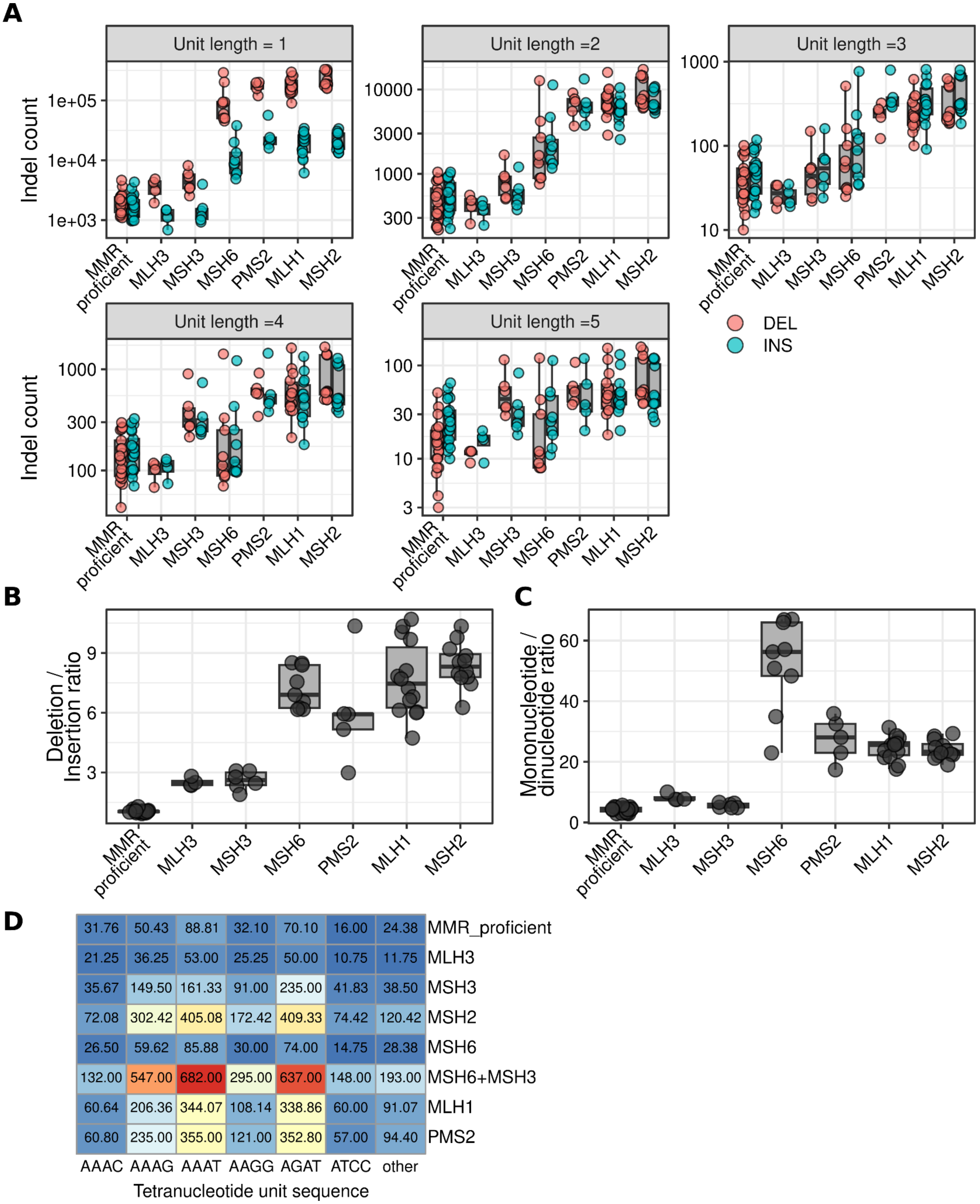


**Supplementary figure S16. Descriptive statistics of short tandem repeat indels in ourK562 cell line panel WGS. A.** Each subpanel shows the absolute deletion and insertion numbers of each MMR genotype, grouped by repeat unit lengths. **B.** Deletion / insertion ratios of MMR mutant genotypes. **C.** Mononucleotide/dinucleotide ratios across K562 MMR mutant genotypes. D.

Numbers of indels at tetranucelotide repeats per genotype, grouped by the sequence of the repeat unit. Only the six most commonly mutated units are shown. The genotype MSH6-MSH3 is also shown separately to illustrate the phenocopy of the MSH2 genotype.

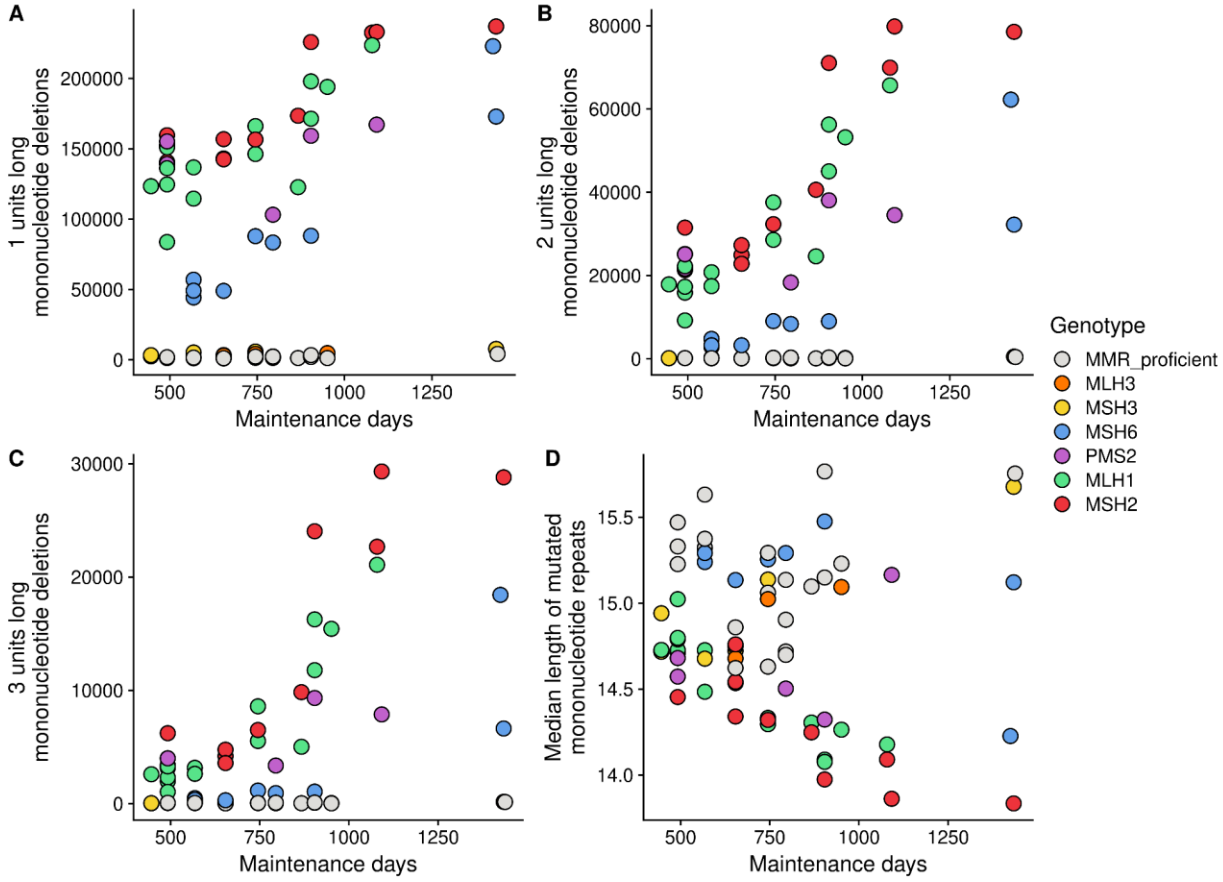


**Supplementary figure S17. Association of mutation accumulation time with STR mutagenesis patterns. A-C.** 1, 2, and 3 unit long deletion numbers at mononucleotide repeats in MSH2, MSH6, MLH1 and PMS2 mutant K562 cells show a positive relationship between time and deletion numbers, with increasing time-dependent rates. **D.** The median length of mononucleotide repeats with deletions are inversely associated with maintenance time, suggesting that with time, shorter repeats start mutating as well.

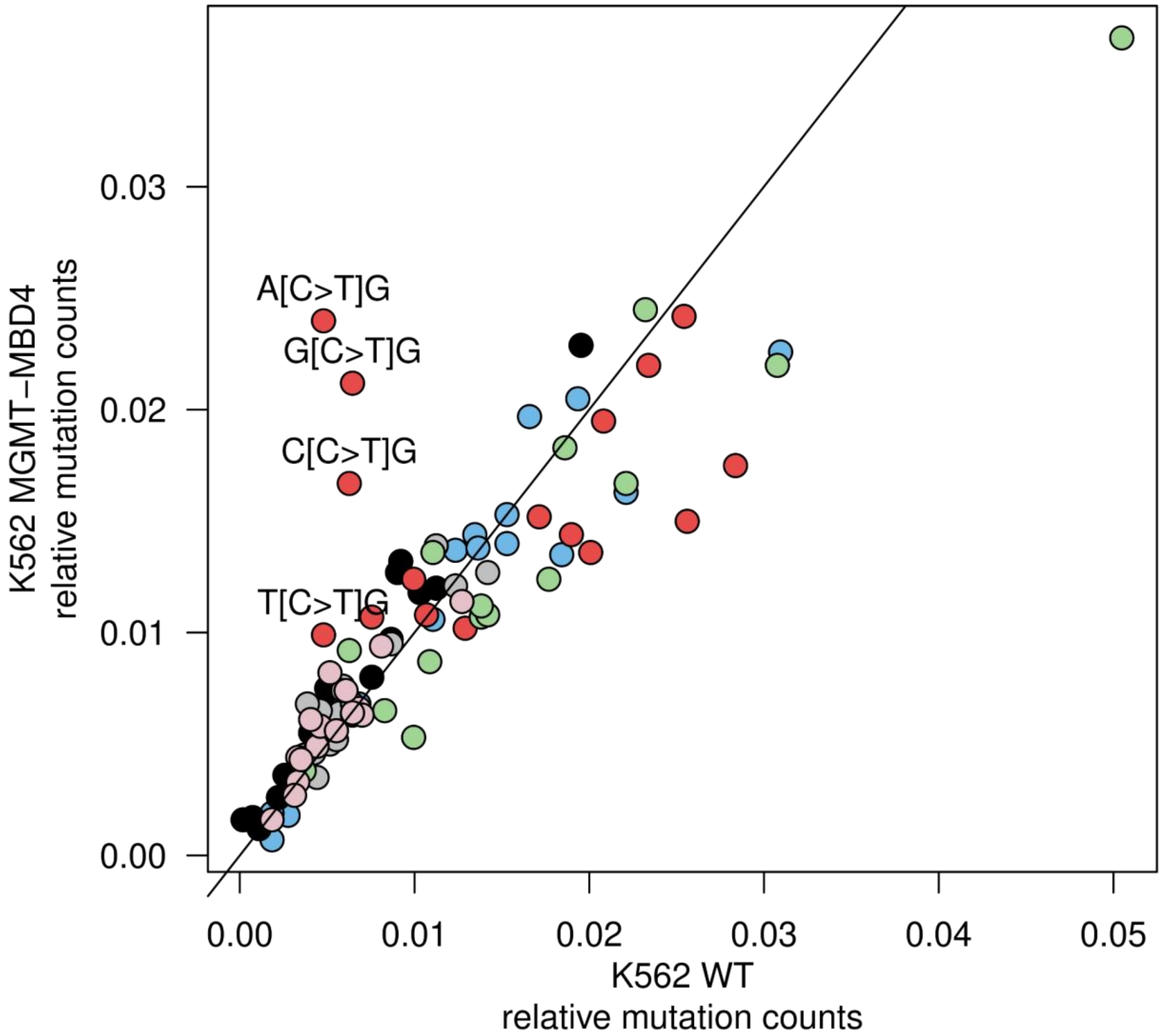


**Supplementary figure S18. Relative mutation frequencies in SBS96 SNV spectra in WT vs MGMT-MBD4 double mutant K562 samples.**
